# BRCA1/BARD1 intrinsically disordered regions facilitate chromatin recruitment and ubiquitylation

**DOI:** 10.1101/2022.08.09.503292

**Authors:** Samuel R. Witus, Lisa M. Tuttle, Wenjing Li, Alex Zelter, Meiling Wang, Klaiten E. Kermoade, Damien B. Wilburn, Trisha N. Davis, Peter S. Brzovic, Weixing Zhao, Rachel E. Klevit

## Abstract

BRCA1/BARD1 is a tumor suppressor E3 ubiquitin (Ub) ligase with roles in DNA damage repair and in transcriptional regulation. BRCA1/BARD1 RING domains interact with nucleosomes to facilitate mono-ubiquitylation of distinct residues on the C-terminal tail of histone H2A. These enzymatic domains constitute a small fraction of the heterodimer, raising the possibility of functional chromatin interactions involving other regions such as the BARD1 C-terminal domains that bind nucleosomes containing the DNA damage signal H2A K15-Ub and H4 K20me0, or portions of the expansive intrinsically disordered regions found in both subunits. Herein, we reveal novel interactions that support robust H2A ubiquitylation activity mediated through a high-affinity, intrinsically disordered DNA-binding region of BARD1. These interactions support BRCA1/BARD1 recruitment to chromatin and sites of DNA damage in cells and contribute to their survival. We also reveal distinct BRCA1/BARD1 complexes that depend on the presence of H2A K15-Ub, including a complex where a single BARD1 subunit spans adjacent nucleosome units. Our findings identify an extensive network of multivalent BARD1- nucleosome interactions that serve as a platform for BRCA1/BARD1-associated functions on chromatin.

## Introduction

Mutations in *BRCA1* and *BARD1* increase lifetime risk of breast and ovarian cancer. BRCA1 and BARD1 form a large, obligate heterodimeric complex (BRCA1/BARD1) that has distinct roles in DNA double-stranded break (DSB) repair by homologous recombination (HR), transcriptional regulation, and several other nuclear processes (Densham & Morris, 2019; Mullan et al., 2006; Tarsounas & Sung, 2020). These functions are mediated, at least in part, through a direct association with chromatin. Upon DNA damage, BRCA1/BARD1 is recruited to damaged chromatin where it segregates with DNA repair factors (Kolas Nadine K. et al., 2007; Mattiroli et al., 2012; Scully et al., 1997; Sobhian Bijan et al., 2007). Cancer-predisposing mutations that disrupt BRCA1/BARD1 functions on chromatin lead to DNA-damage hypersensitivity and genomic instability (Alenezi et al., 2020; Gudmundsdottir & Ashworth, 2006; Krais & Johnson, 2020). BRCA1/BARD1 also has distinct roles in both stimulation and repression of transcription through as yet incompletely defined mechanisms (Mullan et al., 2006). Full knowledge of how BRCA1/BARD1 interfaces with chromatin, and where it exerts its major biological functions, is critical for understanding the etiology of BRCA1/BARD1-mutant associated breast and ovarian cancer.

Nucleosomes, the organizing unit of chromatin, directly interact with BRCA1/BARD1 through multiple direct binding interfaces (Witus et al., 2022). The N-terminal RING domains of BRCA1 and BARD1 form a heterodimer that constitute a RING-type E3 ubiquitin (Ub) ligase and provide the only known enzymatic activity of BRCA1/BARD1 (Brzovic et al., 2001; Lorick et al., 1999; Witus, Stewart, et al., 2021). The RING heterodimer binds to the histone surface of one pseudo-symmetrical ‘face’ of a nucleosome to facilitate site-specific transfer of mono-Ub to a cluster of lysine residues on the extreme C-terminal tail of canonical histone H2A variants (K125, K127, and K129; collectively referred to henceforth as K127) (Hu et al., 2021; Kalb et al., 2014; Witus, Burrell, et al., 2021). H2A lysine specificity is largely determined by the RING domain of BARD1.

This domain binds to a unique histone surface compared to structurally similar Ub ligase complexes that target other H2A lysine residues (e.g., K119 by RING1B/BMI1) (McGinty et al., 2014). The H2A ubiquitylation (“H2A-Ub”) activity of BRCA1/BARD1 is thought to promote long-range resection of broken DNA ends during DSB repair via HR (Densham et al., 2016; Uckelmann et al., 2018). BRCA1/BARD1-dependent H2A-Ub activity is also implicated in the transcriptional repression of certain estrogen-metabolizing cytochrome P450 genes and the constitutive repression of α-satellite DNA regions (Stewart et al., 2018; Thapa et al., 2022; Q. Zhu et al., 2011, 2018).

The BARD1 C-terminal domains (CTDs; **Figure 1A**) also bind to nucleosomes to facilitate DNA DSB repair (Becker et al., 2021; Dai et al., 2021; Hu et al., 2021; Krais et al., 2021; Nakamura et al., 2019; Sherker et al., 2021). The BARD1 CTDs specifically recognize nucleosomes that contain H2A K15-Ub (the product of the Ub ligase RNF168) and unmethylated H4 K20 (H4K20me0). Together, these signals serve as a binding platform for the recognition of damaged chromatin in S/G2 phases when a newly replicated sister chromatid is available to template high-fidelity DNA repair via HR. The BARD1 CTDs and N-terminal BRCA1/BARD1 RING domains bind to histone surfaces that overlap, precluding their simultaneous binding to one nucleosome face. Despite this, mono-nucleosomes with ubiquitin pre-installed at both copies of H2A K15 residues are better substrates for BRCA1/BARD1-dependent ubiquitylation of H2A K127 *in vitro* (Hu et al., 2021). This observation implies that both domains may bind to the same nucleosome unit, with each occupying one face. Additionally, regions throughout the expansive intrinsically disordered segments of both BRCA1 and BARD1 are reported to bind to DNA, but the functional significance of such interactions in chromatin binding and H2A-Ub activity remains to be determined (Mark et al., 2005; Masuda et al., 2016; Simons et al., 2006; Zhao et al., 2017).

**Figure 1.**
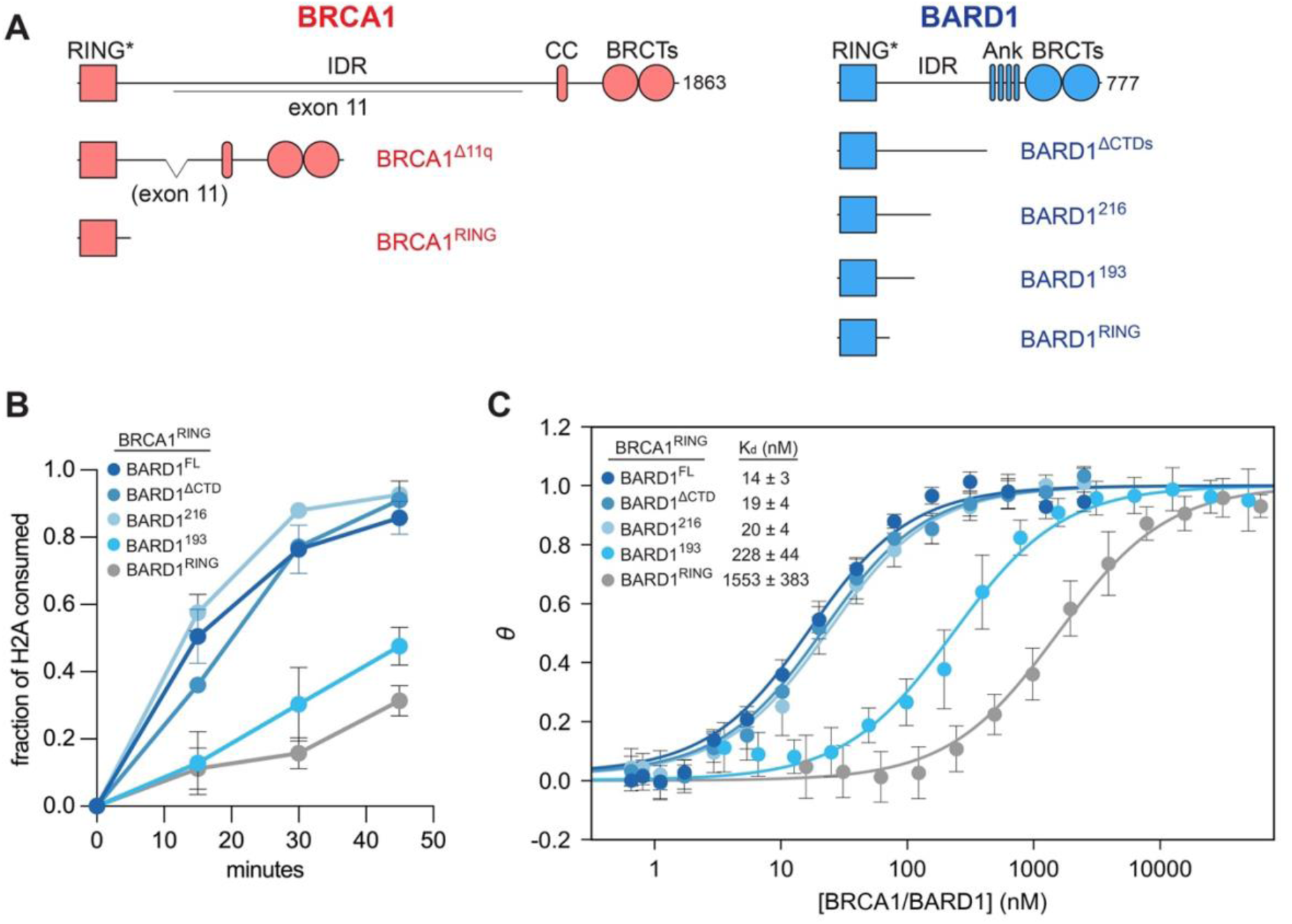
Contributions of BARD1 regions to nucleosome binding and H2A-Ub activity. **A.** Domain organization of BRCA1 and BARD1. The domain names are indicated above the cartoons (RING, really interesting new gene; IDR, intrinsically disordered region; CC, coiled-coil; BRCT, BRCA1 C-terminal; Ank, Ankyrin repeat domain; CTD, C-terminal domain). Constructs used to generate data in panels b and c are shown, and residue bounds are reported in Extended Data Figure I. **B.** Quantification of time-course nucleosome H2A-Ub assays using the indicated BRCA1/BARD1 truncations. Data points represent mean values and error bars are ± 1-s.d. of n=3 independent experiments. **C.** Comparison of nucleosome binding affinity from fluorescence-quenching based measurements using the indicated BRCA1/BARD1 constructs. Data show mean values and error bars are ± 1-s.d. of n=4 independent experiments. The reported difference in affinities to BRCA1^RING^/BARD1^RING^ is likely underestimated, as the minimal RING/RING heterodimer binding data was collected at a lower ionic strength than the rest of the constructs to obtain well-behaved binding data (50 mM vs. 100 mM NaCl).

Although a minimal RING/RING heterodimer is sufficient to direct the transfer of mono-Ub to H2A K127 in nucleosomes, full-length BRCA1/BARD1 heterodimers form stronger complexes with unmodified nucleosome substrates and exhibit increased H2A-Ub activity (Hu et al., 2021; Witus, Burrell, et al., 2021). This suggests that additional interactions between full-length BRCA1/BARD1 and nucleosomes can occur in the absence of the DNA-damage specific H2A K15-Ub histone marks. Because BRCA1/BARD1 performs diverse biological functions in the nucleus including, but not limited to, DNA repair via HR, transcriptional regulation, stalled replication fork protection, centrosome regulation, R-loop resolution, and cell-cycle regulation, it is likely that the complex can be recruited to chromatin via signals not limited to H2A K15-Ub. In pursuit of a wholistic understanding of BRCA1/BARD1 recruitment to chromatin, we sought to characterize interactions with nucleosomes that lead to enhanced chromatin binding and H2A- Ub activity in both the absence and presence of H2A K15-Ub marks.

Here, we identify an intrinsically disordered region of BARD1 adjacent to its RING domain that binds strongly to both nucleosomal and extra-nucleosomal DNA, dramatically enhancing the affinity of the complex and its H2A-Ub activity. The interactions can be modulated by specialized DNA structures that are derived from the DNA damage repair process and can compete directly with BARD1 binding to nucleosomal DNA. We incorporate our findings into a molecular mechanism of recognition and establish a role for BARD1-DNA interactions in chromatin recruitment and DNA damage repair in cells. Additionally, we provide evidence for multiple, distinct higher-order chromatin complexes that contain H2A K15-Ub nucleosomes. These include both a ‘wrapped’ complex with the BRCA1/BARD1 RING domains and BARD1 CTDs bound to opposite sides of one nucleosome unit, as well as an extended complex where the domains bind to adjacent nucleosome units. Throughout, we evaluate the contribution of BARD1 DNA binding to nucleosome affinity and H2A-Ub activity. Our findings reveal a network of multivalent BARD1-nucleosome interactions that serve as a platform for BRCA1/BARD1-associated functions on chromatin and hint at novel modes of DNA recognition by intrinsically disordered regions of proteins.

## Results

### BARD1 IDR supports increased H2A ubiquitylation activity

The BRCA1/BARD1 heterodimer contains both structured domains and long intrinsically disordered regions (IDRs**; Figure 1A)**. A minimal complex composed of the N-terminal RINGs of BRCA1 (1-112) and BARD1 (26-140; BRCA1^RING^/BARD1^RING^) is sufficient to direct site-specific mono-ubiquitylation of nucleosomal histone H2A at K127 (Kalb et al., 2014). We and others have observed that full-length BRCA1/BARD1 (∼300 kDa) exhibits faster H2A-Ub kinetics and stronger binding to unmodified mono-nucleosomes than the minimal RING/RING complex (∼25 kDa; **Supplemental Figure 1A-D**) (Hu et al., 2021; Witus, Burrell, et al., 2021). To identify regions responsible for increased H2A-Ub activity and binding affinity, we generated heterodimers with truncations in either BRCA1 or BARD1 **(Supplemental Figure 2 and Extended Data Table I)**.

Heterodimers containing full-length BARD1 and three BRCA1 constructs were compared to survey BRCA1 regions that may enhance H2A-Ub activity: full-length BRCA1 (BRCA1^FL^), a clinically relevant allele missing ∼1100 residues in the expansive IDR (BRCA1^Δ11q^), and the minimal BRCA1 RING fragment (BRCA1^RING^). Notably, in the context of BARD1^FL^, the RING domain of BRCA1 is sufficient to generate fast H2A-Ub kinetics **(Supplemental Figure 1C)**. The presence of additional regions of BRCA1 do not further enhance activity (as in BRCA1^Δ11q^/BARD1^FL^) or appear to inhibit activity (e.g., BRCA1^FL^/BARD1^FL^). These results indicate that the BARD1 subunit is largely responsible for enhanced H2A-Ub activity. In the context of heterodimers that contain only the RING of BARD1, modest activity enhancements are afforded by inclusion of additional BRCA1 regions, suggesting that, in the absence of FL- BARD1, regions of BRCA1 may provide some redundancy in function **(Supplemental Figure 1D)**.

To identify regions of BARD1 that contribute to increased H2A-Ub activity, the H2A-Ub kinetics and nucleosome binding affinity of BRCA1^RING^-heterodimers containing BARD1 truncations were compared. Notably, BRCA1^RING^/BARD1^FL^ heterodimers bind nucleosomes with ∼100-fold higher affinity than minimal BRCA1^RING^/BARD1^RING^ and have markedly higher H2A-Ub activity **(Figure 1B, C)**. Removal of the BARD1 C-terminal domains (BARD1^ΔCTD^) or truncation back to residue 216 in the putative intrinsically disordered region (BARD1^216^) had minimal impact on H2A-Ub activity and nucleosome binding **(Figure 1B, C)**. However, truncation to residue 193 (BARD1^193^) caused a substantial decrease in H2A-Ub activity and a ∼10-fold reduction in binding affinity, suggesting a nucleosome-binding interface for residues between 194 and 216. Consistent with this, deletion of the first half of the BARD1 IDR (BARD1^Δ140-270^) greatly diminished H2A-Ub activity **(Supplemental Figure 1E)**. Although deletion of residues 140-270 decreased nucleosome binding affinity by ∼3-fold relative to full-length BARD1, the affinity is still considerably stronger than the minimal RING/RING heterodimer or BARD1^193^ truncations **(Supplemental Figure 1F)**. A more targeted deletion (BARD1^Δ194-216^) only modestly decreased H2A-Ub activity, suggesting the presence of additional or compensatory nucleosome interaction sites in BARD1 *in vitro*. Intrinsic nucleosome-binding properties of BARD1 IDRs were analyzed using constructs containing only the N-terminal half (residues 124-270) or only the C-terminal half (residues 269-424). Each IDR-only construct exhibited nucleosome binding by EMSA, with modest differences in affinity **(Supplemental Figure 1G).** In summary, only the RING domain of BRCA1 is required for high H2A-Ub activity in vitro while a small region within the BARD1 IDR (BARD1 141-216, referred to in the discussion as IDR^Prox^) that engages in nucleosome binding is required in addition to the BARD1 RING to support increased H2A-Ub activity.

### Characterization of the BARD1 IDR reveals that the additional BARD1-nucleosome interactions are mediated through DNA binding

BARD1 residues 141-216 identified as contributing to increased H2A-Ub activity and nucleosome binding are in the large central segment predicted to be intrinsically disordered (124-424) (McGuffin et al., 2000). Nuclear magnetic resonance (NMR) ^1^H^15^N-HSQC spectra of ^15^N-BARD1 fragments containing residues 124-270, 269-424, and 124-424 each exhibited narrow ^1^H chemical shift dispersion consistent with intrinsic disorder **(Supplemental Figure 3A, B)**. Furthermore, individual spectra of BARD1 IDRs 124-270 and 269-424 overlay well with the spectrum of the full region (124-424), indicating that the two regions behave independently of one another **(Supplemental Figure 3B)**.

Despite being intrinsically disordered, the BARD1 124-270 sequence is largely conserved across orthologs **(Supplemental Figure 3C)**. The region was previously reported to bind to double-stranded DNA and specialized DNA structures containing base-pair mismatch (discussed below) (Zhao et al., 2017). We therefore hypothesized that the contributions of the BARD1 IDR (residues 141-216) to increased binding and activity are mediated through interactions with nucleosomal DNA. Indeed, BARD1 124-270 binds strongly to nucleosomes and to free 147-bp ‘601’ dsDNA by EMSA **(Supplemental Figure 4A)**.

For residue-level information, we performed NMR binding experiments with ^15^N-labelled BARD1 124-270 and either a 36-bp dsDNA fragment or mono-nucleosomes wrapped with 147-bp dsDNA **(Figure 2A and Supplemental Figure 4B)**. A full titration was possible only with the dsDNA fragment due to its lower molecular weight and higher solubility. The spectral series displayed a subset of resonances that shift continuously as a function of DNA concentration **(Figure 2A**). A similar set of resonances was perturbed (broadened in this case) upon addition of 147-bp nucleosome, indicating that similar BARD1 residues engage nucleosomes and free dsDNA **(Supplemental Figure 4B)**. To overcome spectral overlap in the spectrum of BARD1 124-270, the spectrum of a smaller construct containing the putative DNA binding region (BARD1 141-216, where residue 141 is the first to extend beyond BARD1^RING^) was assigned and assignments were transferred to the longer segment, enabling analysis in both contexts **(Figure 2A, B and Supplemental Figure 4B, C)**. Secondary structure propensity (SSP) predicted from the resonance assignments are consistent with the region being primarily disordered, with some helical propensity predicted for residues 191-201 and a weaker prediction of extended structure towards the beginning of the construct **(Supplemental Figure 5A).**

**Figure 2.**
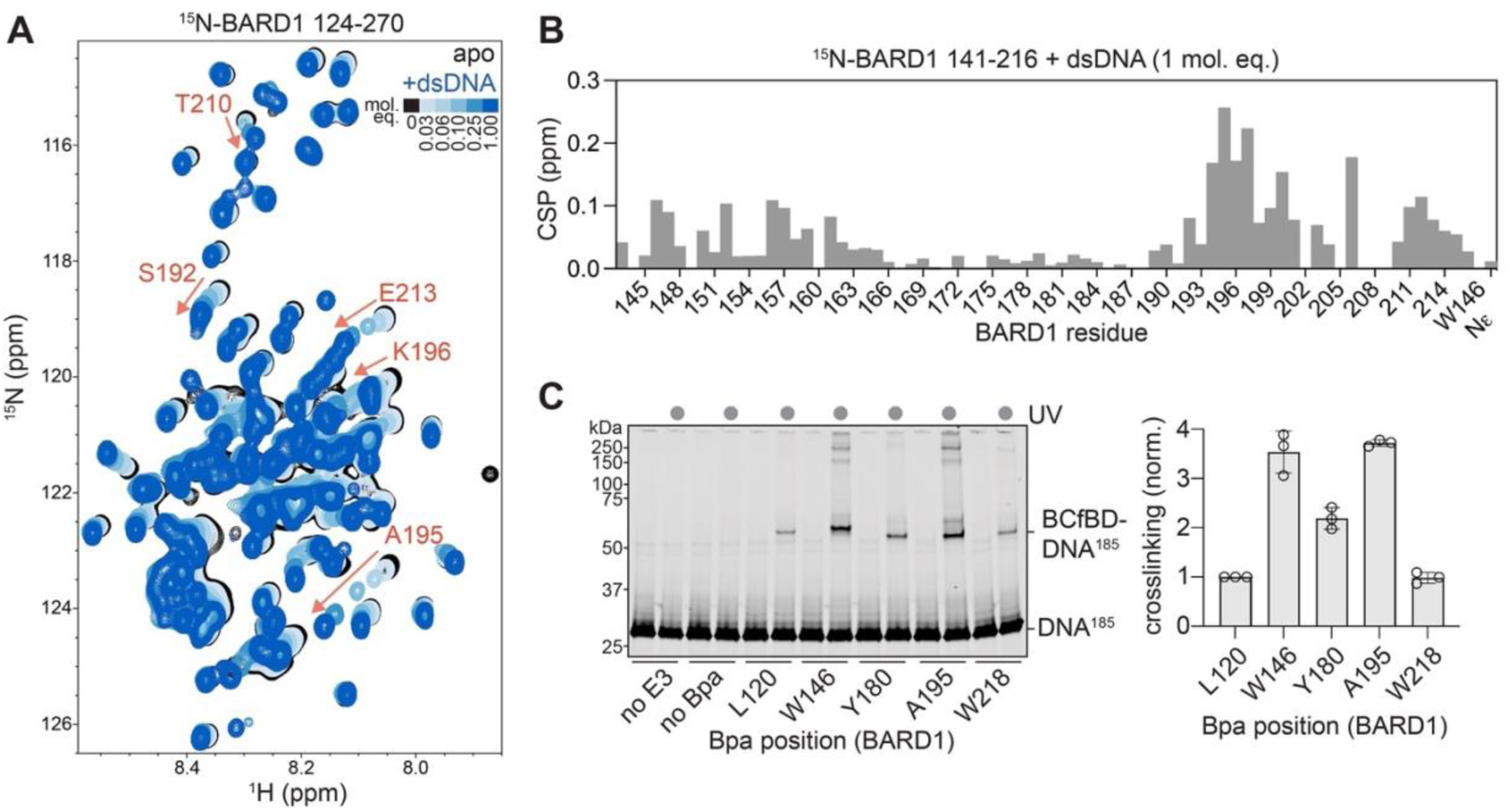
The BARD1 IDR binds to nucleosomes via DNA. **A.** ^1^H^15^N-HSQC NMR titration spectra of ^15^N-BARD1 124-270 with a 36-bp dsDNA fragment. Darker shades of blue correspond to increasing amounts of DNA added. A subset of highly affected signals is labelled with their residue identities, with arrows showing their trajectories over the course of the titration. **B.** ^1^H^15^N-HSQC NMR amide chemical shift perturbations (CSPs) observed to ^15^N-BARD1 141- 216 signals when bound to a 36-bp dsDNA fragment (1:1 molar equivalent complex). **C.** Representative SDS-PAGE gel monitoring in-gel fluorescence (labelled DNA) of UV-induced Bpa crosslinking of BRCA1-f-BARD1^221^ to nucleosomes (left). Quantification of Bpa crosslinking experiments with nucleosomes (right). The intensity of each crosslinked band was normalized to the intensity of the L120Bpa crosslinked band for each replicate experiment. Graph bars show the mean; error bars are ± 1-s.d. and the open circles are the values of individual replicates for of n=3 independent experiments.

BARD1 residues 194-216 exhibit the largest backbone amide chemical shift perturbations (CSPs) due to dsDNA addition, with smaller perturbations observed for residues 146-161 **(Figure 2B).** Residues between 162-191 were relatively unaffected by dsDNA, consistent with the existence of distinct DNA-binding regions within the IDR. Secondary structure predicted from chemical shifts in the presence of dsDNA (1 molar equivalent) indicate reduced helical propensity in residues 191-199 upon DNA binding **(Supplemental Figure 5A)**. As well, ^15^N dynamics measurements reveal that residues 146-161 undergo changes in dynamics in the presence of dsDNA **(Supplemental Figure 5B)**. Although low solubility limited similar analysis on nucleosome-bound samples, the similar spectral perturbations observed in the presence of low molar equivalents of nucleosomes or dsDNA suggest similar protein-DNA binding modes **(Supplemental Figure 4B)**. Thus, the data reveal two IDR regions adjacent to the RING domain (146-161 and 194-216) that bind DNA when presented as free dsDNA or in the context of a nucleosome. These regions are consistent with the H2A-Ub activity and nucleosome binding results presented above.

Having established that isolated BARD1 IDR fragments bind to nucleosomes, we sought to observe the interactions within the context of an active enzymatic complex. As such a complex is too large for NMR analysis, we employed targeted crosslinking of BRCA1/BARD1 to nucleosomal DNA. Photoactivatable crosslinker p-Benzoyl-L-phenylalanine (Bpa) was incorporated at individual positions in the BARD1 IDR in the context of RING-containing constructs using amber-codon suppression in *E. coli* (L120Bpa, W146Bpa, Y180Bpa, A195Bpa and W218Bpa**; Supplemental Figure 6A)**. Due to the need to co-express a Bpa-specific aminoacyl tRNA transferase, we designed a genetic fusion in which BRCA1 residues 1-104 are fused to BARD1 residues 26-221 through a 12-residue GlySer-linker (BRCA1-f-BARD1^221^).

Wild-type BRCA1-f-BARD1^221^ and Bpa-incorporated mutants retained similar H2A-Ub activity to each other and to unfused heterodimers **(Supplemental Figure 6B)**. Nucleosomes with extra- nucleosomal linker DNA that carried a fluorescent probe enabled visualization of covalent BARD1-DNA crosslinked products via SDS-PAGE (NCP^185^; **Supplemental Figure 6C)**. Such bands were detected for every Bpa variant, with enhanced crosslinking by variants with Bpa at BARD1 W146 and A195 positions, consistent with the NMR dsDNA binding data presented above **(Figure 2C)**. Crosslink bands were weaker from Y180 and W218 positions and comparable to crosslinks from L120 which is located close to the RING domains and not expected to contribute to productive DNA interactions. A similar pattern was observed in crosslinking reactions containing free DNA **(Supplemental Figure 6D)**. As crosslinks can only form from Bpa to sites that are within ∼3Å, the data support a direct interaction between the BARD1 IDR and nucleosomal DNA in the context of an enzymatically active complex.

### Contribution of DNA binding to enhanced H2A-Ub activity

Results presented thus far establish that the DNA-binding region of the BARD1 IDR is necessary for high H2A-Ub activity, but is it sufficient? To address this question, we asked whether the DNA-binding properties of full-length BARD1 can support H2A-Ub activity in the presence of BARD1 RING mutations that disrupt the histone-binding interface (BARD1 P89A/W91A; **Figure 3A)** (Hu et al., 2021; Witus, Burrell, et al., 2021). Heterodimers containing BRCA1^RING^ and full-length BARD1 (P89A/W91A) retain high-affinity nucleosome binding and auto-ubiquitylation activity but do not transfer Ub to H2A or any other histone sites **(Figure 3B, Supplemental Figure 7A-C)**. Thus, the BARD1 RING-histone interface is absolutely required for H2A-Ub activity while the BARD1 IDR-DNA interactions promote formation of a high-affinity complex that supports enhanced H2A-Ub activity.

**Figure 3.**
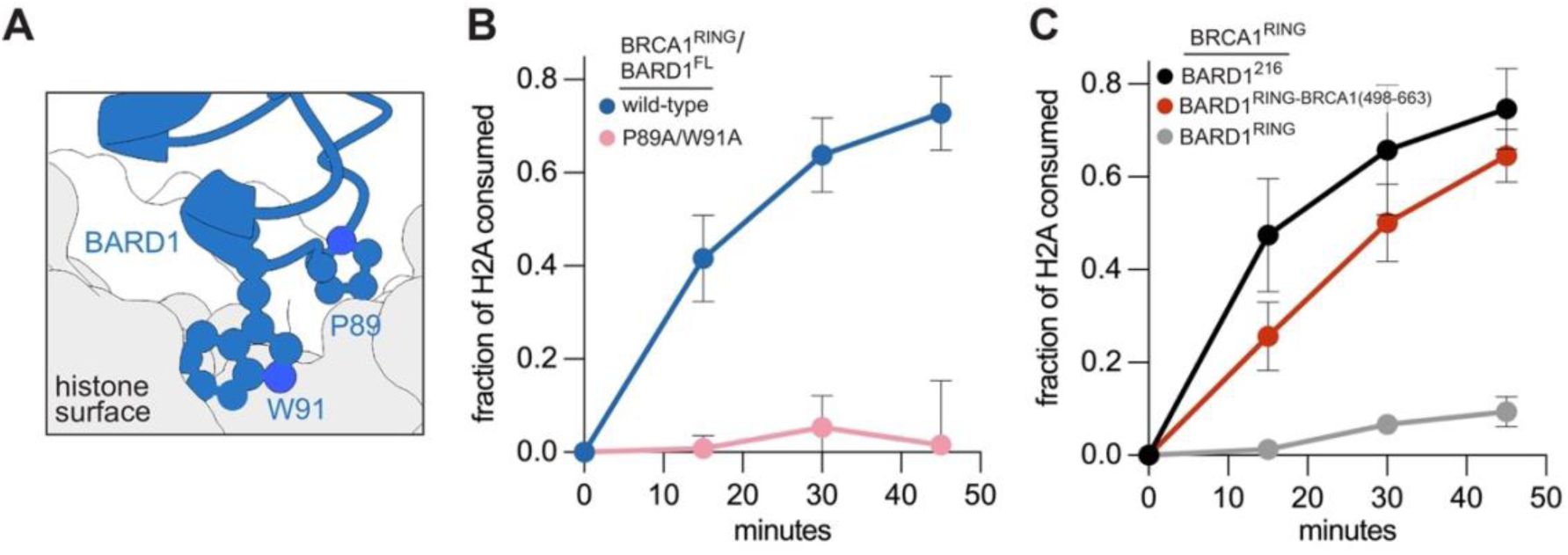
Contribution of BARD1 DNA-binding and RING-histone binding to H2A-Ub activity. **A.** The BARD1 RING-histone interface showing the locations of BARD1 P89 and W91 relative to the histone surface (PDB: 7JZV). **B, C.** Quantification of time-course H2A-Ub assays using the indicated BRCA1/BARD1 constructs. Data show the mean; error bars are ± 1-s.d. of n=3 independent experiments. Two single time-points were determined to be outliers in panel C and excluded from analysis. Values for these excluded points are included as source data.

The BARD1 DNA-binding region(s) are disordered. To test whether any DNA-binding region placed adjacent to the RING domain can serve the same purpose, we created a chimeric BARD1 construct in which a previously identified DNA-binding region from the BRCA1 IDR (498-663) was appended onto the BARD1 RING (Mark et al., 2005). A heterodimer composed of BRCA1^RING^ and the BARD1 chimera displayed increased H2A-Ub activity compared to the minimal RING/RING complex, but not to the level of the native BARD1 sequence **(Figure 3C)**. The nucleosome-binding affinities for these species are quite similar, suggesting that the modest activity difference between the native BARD1 region and the installed BRCA1-derived region arises from additional features **(Supplemental Figure 7D)**. The findings indicate that a major contributor to increased H2A-Ub activity is the presence of a disordered, high-affinity DNA-binding region near the enzymatic RING domains, but indicate that the BARD1 IDR is especially effective.

### Specialized DNA structures bind to the BARD1 IDR and inhibit H2A-Ub activity

Based on a previous report that the BARD1 IDR preferentially binds to partly unwound DNA structures (bubble-DNA) and homologous recombination (HR) intermediates (D-loop DNA) over double- or single-stranded DNA (Zhao et al., 2017), we hypothesized that interaction of the BARD1 DNA-binding region with bubble- or D-loop DNA might affect its ability to interact with nucleosomal DNA. Bubble- and D-loop- DNA substrates are likely similar based on the near-identical affinities of the BARD1-DNA complexes (Zhao et al., 2017), so we used the simpler bubble-DNA substrate in our assays. In nucleosome ubiquitylation reactions carried out with full-length BRCA1/BARD1 in the presence of equal concentrations of DNA fragments containing various patterns of base-pair mismatch, only bubble-DNA substantially inhibited H2A-Ub activity. **(Figure 4A, B and Supplemental Figure 8A, B)**. The minimal RING heterodimer activity was unaffected by bubble-DNA, consistent with the inhibition being due to competing interactions of bubble- and nucleosomal DNA for binding to BARD1 141-216, **(Supplemental Figure 8C, D)**. BARD1-nucleosome DNA crosslink formation was inhibited by either bubble-DNA or dsDNA inhibited in a concentration-dependent manner, with bubble-DNA being substantially more effective **(Figure 4C and Supplemental Figure 9).**

**Figure 4.**
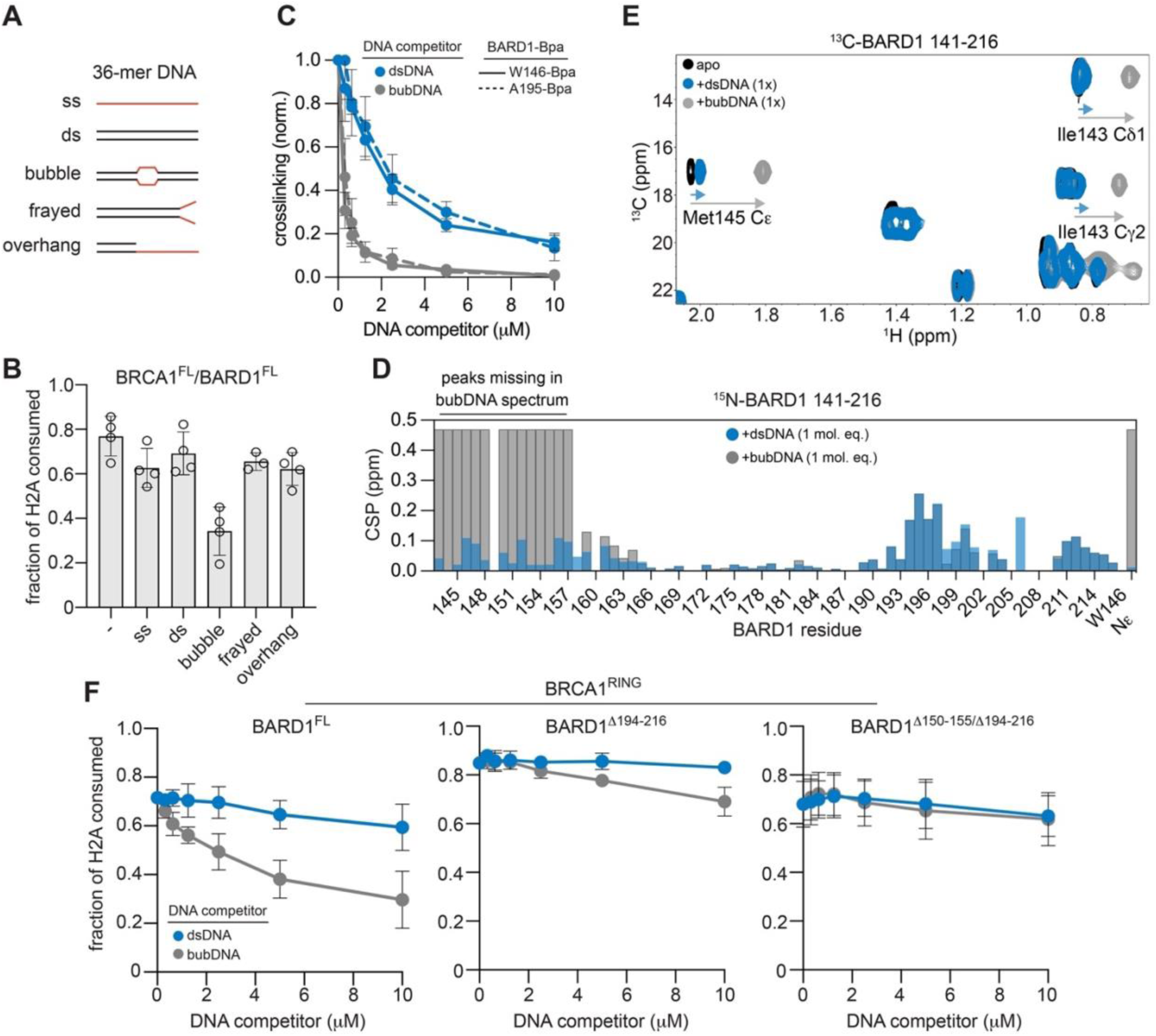
Inhibition of nucleosome binding and H2A-Ub by specialized DNA structures. **A.** Design of 36-mer DNA competitor fragments used in H2A-Ub activity, Bpa crosslinking, and NMR assays. Red indicates non-base-paired regions. The number of unpaired bases or base- pairs is: ss – 36; ds – 0; bubble – 8; frayed – 8; ssOverhang – 24. **B.** Single time-point H2A-Ub inhibition assays using BRCA1^FL^/BARD1^FL^ (100 nM) and the indicated competitor DNA (2.5 µM). Data show the mean; error bars are ± 1-s.d.; and open circles are individual data points of n=4 independent experiments. One outlier point for “frayed- DNA” was excluded from the analysis. **C.** Inhibition of UV-induced Bpa crosslinking between the indicated BRCA1-f-BARD1^221^ Bpa- incorporated constructs and NCP^185^ substrates in the presence of increasing amounts of dsDNA or bubble-DNA competitor. Data show the mean; error bars are ± 1-s.d. of n=3 (W146Bpa) or n=4 (A195Bpa) independent experiments. **D.** ^1^H^15^N-HSQC NMR CSPs observed to ^15^N-BARD1 141-216 signals when bound to a 36-mer dsDNA (blue, front) or bubble-DNA (gray, back) fragment (1:1 molar equivalent complex). Signals for residues 143-157 are broadened beyond detection in the bubble-DNA-bound spectrum, and gray bars are equal to the CSP value for the W146Ne CSP that was observed (see Supplemental Figure 10A, B for spectra). **E.** Selected region of ^13^C-HSQC spectra of ^13^C-BARD1 141-216 in 1:1 molar equivalent complex with dsDNA fragment or bubble-DNA fragment. Signal trajectories in bound spectra are indicated by arrows. **F.** Single time-point H2A-Ub inhibition assays using heterodimers containing BRCA1^RING^/BARD1^FL^ (left, 12 min endpoint) or the indicated BARD1 internal deletion mutant (middle and right, 20 min endpoint) and increasing amounts of dsDNA or bubble-DNA competitor. The same amount of E3 (50 nM) was used for each BRCA1/BARD1 truncation. Data show the mean; error bars are ± 1-s.d. of n=3 independent experiments. One time-point from Panels B and C each were determined to be outliers and excluded from analysis. Values for these points are included as source data.

We compared the interactions of BARD1 141-216 with bubble- and dsDNA by NMR using DNA species of similar sequence and size. CSPs and changes in dynamics were similar within BARD1 194-216, the region identified as the major site for dsDNA and nucleosomal DNA binding **(Figure 4D and Supplemental Figure 10A-C)**. Much stronger effects were produced by bubble-DNA on backbone resonances of residues 143-157 with severe peak broadening/disappearance, indicative of large changes in chemical environment and/or dynamics. Only in the presence of bubble-DNA was a large CSP observed for the sidechain resonance of W146 **(Supplemental Figure 10A, B).** Additional effects on sidechains were observed in ^1^H^13^C-HSQC spectra of BARD1 141-216, where methyl group peaks from BARD1 residues 143-157 exhibit large CSPs that are not observed in the presence of dsDNA **(Figure 4E and Supplemental Figure 10D)**. Together, the data show that bubble DNA engages strongly and specifically at a site near the beginning of the BARD1 IDR (residues 143-157) and that both forms of DNA engage BARD1 194-216. Both IDR regions appear to be functionally important, as only concomitant deletion of basic residues in both IDR regions (BARD1^Δ150-155/Δ194-216^) alleviated H2A-Ub activity inhibition by bubble-DNA **(Figure 4F and Supplemental Figure 8E)**.

### DNA-binding IDR facilitates chromatin recruitment and DNA damage repair in cells

To test the importance of BARD1 IDR DNA binding in a cellular context, we established doxycycline-inducible HeLa cell lines that deplete endogenous BARD1 via doxycycline-induced expression of shRNA against BARD1 (HeLa-shBARD1) and stably re-express shBARD1- resistant HA-tagged BARD1 (wild-type or Δ194-216; **Supplemental Figure 11A**). Endogenous BARD1 is largely depleted (>90%) in these cells and both wild-type BARD1 and BARD1^Δ194-216^ localize to the nucleus and are expressed at similar levels **(Supplemental Figure 11A, B)**. In cellular fractionation assays, less BARD1^Δ194-216^ is associated with chromatin than wild-type BARD1 under both basal and olaparib-induced DNA-damage conditions **(Figure 5A)**.

**Figure 5.**
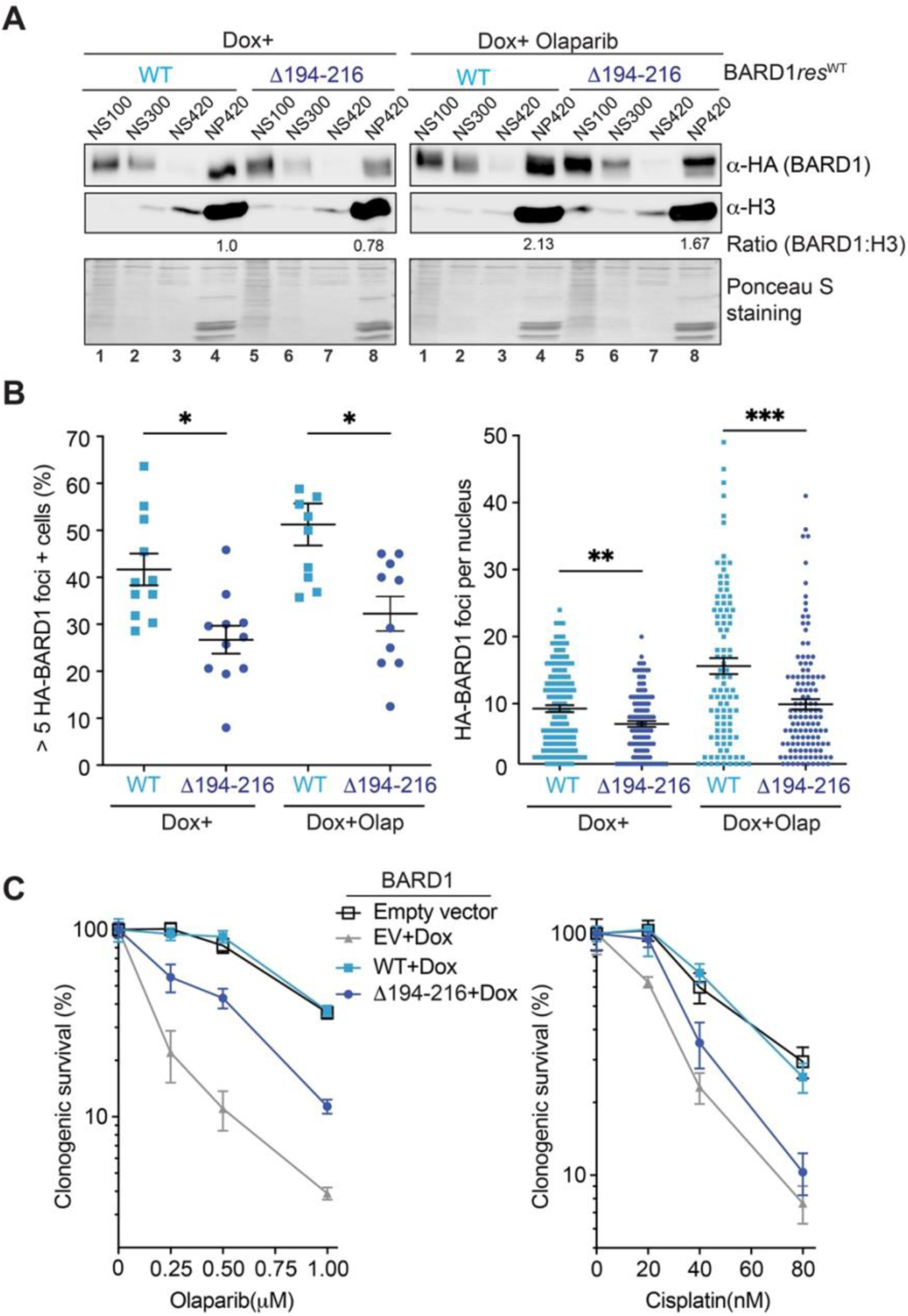
Cellular role for BARD1 DNA-binding in chromatin recruitment and DNA damage repair. **A.** Western blot analysis to detect HA-BARD1 in nuclei isolated from HeLa-shBARD1 cells stably expressing wild type or Δ194-216 mutant of HA-BARD1, where endogenous BARD1 was depleted by doxycycline-induced shBARD1 expression. The nuclei were salt-fractionated into NS100, NS300, NS420, and NP420 to assess the amount of HA-BARD1 associated with chromatin. **B.** Quantification of cells with >5 HA-BARD1 foci (left panel) and number of HA-BARD1 foci per nucleus (right panel) with and without Olaparib treatment. *p<0.05, **p<0.01, ***p<0.001, by Student’s t-test. **C.** Clonogenic survival of HeLa-shBARD1 cells stably expressing wild type or Δ194-216 mutant of HA-BARD1 upon their treatment with indicated amount of olaparib and cisplatin. Error bars represent SEM (n=3).

Significantly reduced HA-BARD1^Δ194-216^ foci formation was observed in both untreated and olaparib-induced DNA-damage conditions, indicating a defect in BRCA1/BARD1 recruitment to chromatin and DNA damage sites **(Figure 5B and Supplemental Figure 11C)**. To assess the ability of the BARD1 IDR mutant to facilitate DNA break repair, we performed clonogenic survival assays upon treatment with DNA-damaging agents olaparib and cisplatin **(Figure 5C and Supplemental Figure 11D)**. Survival of BARD1^Δ194-216^ cells was significantly reduced compared to wild-type controls under both drug treatments – a hallmark of defective HR **(Figure 5C)**. Altogether, the results demonstrate that the short intrinsically disordered stretch of BARD1 with DNA-binding ability contributes to chromatin association in cells under both basal and exogenous DNA-damage conditions and contributes to DNA break repair required for cell survival.

### Extra-nucleosomal linker DNA enhances H2A ubiquitylation

Chromatin is composed of adjacent nucleosome units separated by linker DNA. The extended domain topology of both BRCA1 and BARD1 with their long IDRs raises the possibility of interactions in higher-order chromatin substrates that are not present in minimal mono- nucleosome substrates. Nucleosome substrates with extra-nucleosomal linker DNA (NCP^185^) and tri-nucleosomes (tri-NCP) were used as substrates in H2A-Ub assays to test for additional chromatin involvement **(Figure 6A and Supplemental Figure 12A)**. In reactions using full-length BRCA1/BARD1, faster H2A-Ub kinetics were observed for NCP^185^ with linker DNA compared to the minimal NCP^147^ substrate **(Figure 6B)**. Notably, no additional increase in H2A- Ub activity was observed for a tri-NCP substrate despite containing a longer linker DNA and multiple histone binding interfaces. The data are consistent with two conclusions: 1) the presence of linker DNA enhances BRCA1/BARD1 H2A-Ub activity and 2) functional interactions are limited to one mono-nucleosome unit, at least in the absence of histone PTMs.

**Figure 6.**
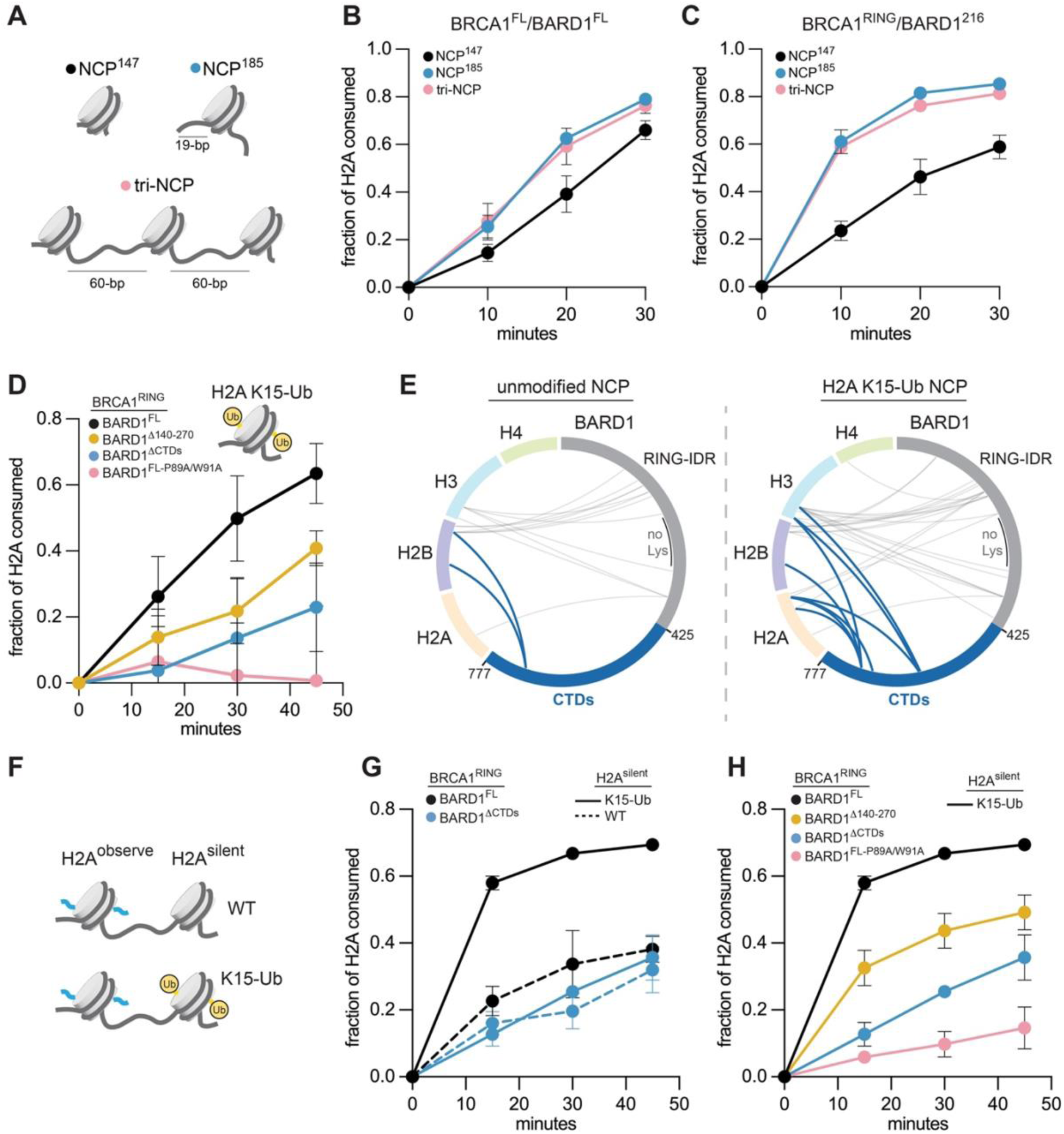
H2A-Ub activity using chromatin substrates with linker DNA and pre-installed H2A K15-Ub. **A.** Schematic of chromatin substrates used for H2A-Ub activity and nucleosome binding assays in panels b and c. The length of linker-DNA is indicated next to the NCP^185^ and tri-NCP. **B, C.** Quantification of time-course H2A-Ub assays using the indicated BRCA1/BARD1 constructs and chromatin substrates. **D.** Quantification of time-course H2A-Ub assays using homogenously modified H2A K15-Ub mono-nucleosome substrates and indicated BRCA1/BARD1 constructs. **E.** Intermolecular crosslinks observed by chemical crosslinking and MS analysis between BARD1 and histones using wild-type (left) and H2A K15-Ub (right) 147-bp ‘601’ nucleosomes and BRCA1^RING^/BARD1^FL^ heterodimers. Crosslinks to histones emanating from the CTDs of BARD1 are shown in blue, and the RING-IDR region in gray. A lysine-depleted region of the BARD1 IDR is labelled and indicated by a black bar. **F.** Design of asymmetric di-NCP substrates. **G, H.** Quantification of time-course H2A-Ub assays using the indicated combinations of di-NCP substrate and BRCA1/BARD1 heterodimer. For panels A, D, and E, data show the mean; error bars are ± 1-s.d. of n=3 independent experiments. One time-point from Panel A was determined to be an outlier and excluded from analysis. The excluded value is reported as source data.

To explicitly test the BARD1 IDR DNA-binding region as the source of the enhanced activity observed for NCP^185^ and tri-NCP substrates, the H2A-Ub activity of the BRCA1^RING^/BARD1^216^ truncation was tested **(Figure 6C)**. This species exhibited considerably faster H2A-Ub kinetics with either NCP^185^ or tri-NCP compared to a minimal NCP^147^ substrate, confirming that the dependence on linker DNA arises from the BARD1 IDR. The presence of linker DNA provides less than a two-fold increase in binding affinity for BRCA1^RING^/BARD1^216^ to NCPs under salt conditions used in H2A ubiquitylation assays **(Supplemental Figure 12B)**. Thus, in the absence of linker DNA, BARD1 IDR can interact nearly as well with nucleosomal DNA, but the interaction with linker DNA supports enhanced activity.

### Effects of H2A-Ub marks on BRCA1/BARD1 H2A-Ub activity

Effects of mono-Ub at H2A positions known to be ubiquitylated by different E3 ligases were assessed using homogenously mono-ubiquitylated nucleosomes containing ubiquitin preinstalled at H2A positions K15, K119, or K127 via a non-hydrolyzable dichloroacetone crosslinker **(Supplemental Figure 13A, B)** (Morgan et al., 2019). Ubiquitin at either H2A C-terminal position (K119 or K127) decreased BRCA1/BARD1-dependent H2A-Ub activity, hinting at exclusivity between BRCA1/BARD1 and RING1/PCGF E3 complexes that both modify the H2A C-terminal tail. Consistent with previous observations, Ub preinstalled at H2A K15 increased the H2A-Ub activity of full-length BRCA1/BARD1, and this enhancement requires BARD1 CTDs **(Figure 6D** and **Supplemental Figure 13C, D)**. This contrasts with unmodified nucleosome substrates where BARD1 CTDs are not required for enhanced H2A-Ub activity (Hu et al., 2021).

### H2A K15-Ub-mediated enhancement of BRCA1/BARD1 H2A-Ub activity

Binding of BARD1 CTDs to nucleosomes that contain both H2A K15-Ub and H4K20me0 (Dai et al., 2021; Hu et al., 2021) recruits BRCA1/BARD1 to damaged chromatin to facilitate DNA DSB repair via HR (Becker et al., 2021; Krais et al., 2021; Nakamura et al., 2019). The BARD1 CTDs use a binding surface that overlaps with the RING domain site, precluding binding of both domains to the same nucleosome face. However, the observation that nucleosomes containing H2A K15-Ub and H4K20me0 are better substrates in the context of BARD1^FL^ implies that RING domains and CTDs can bind simultaneously to opposite faces of a nucleosome (Hu et al., 2021). We used BARD1 truncation constructs to determine what features of BARD1 contribute to enhanced activity on H2A K15-Ub nucleosome substrates. As expected, deletion of the CTDs or histone-binding RING mutations of BARD1 do not support activity **(Figure 6D).** Deletion of the BARD1 DNA-binding IDR (BRCA1^RING^/BARD1^Δ140-270^) led to decreased H2A-Ub kinetics compared with the BRCA1^RING^/BARD1^FL^ complex **(Figure 6D)**. Thus, even with the additional nucleosome-binding contribution provided by the CTDs on H2A K15-Ub nucleosomes, the DNA-binding IDR still plays a role in enhancing ubiquitylation kinetics. Altogether, our findings support a model in which three points of contact between BARD1 and H2A K15-Ub nucleosomes (RING-histone, IDR-DNA, and CTDs-histone/Ub) are required for full BRCA1/BARD1- dependent H2A-Ub activity **(Figure 7E)**.

**Figure 7.**
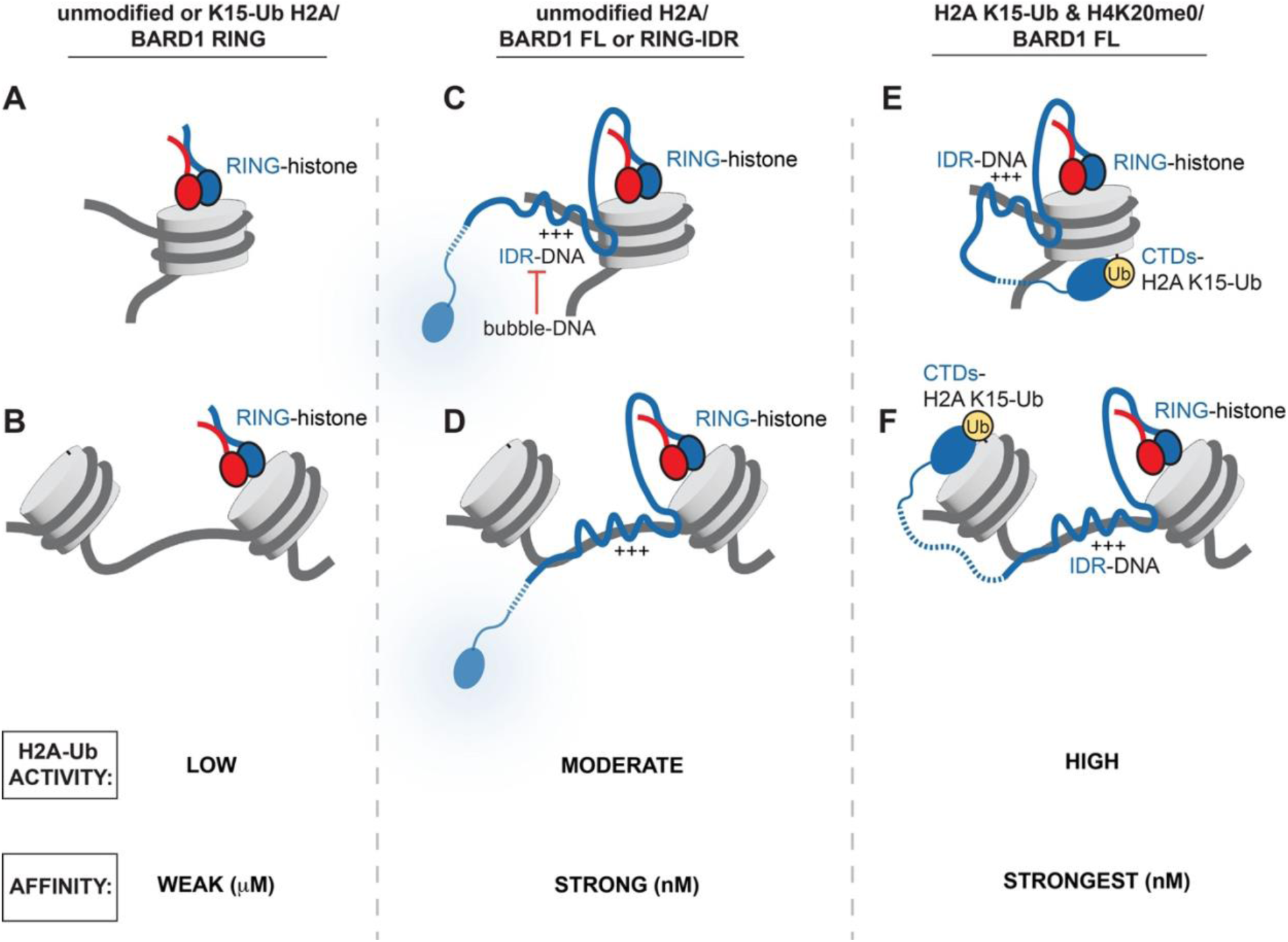
Models of BRCA1/BARD1/chromatin complexes supported by this study. **A, B.** Model of the BRCA1/BARD1 minimal RING heterodimer binding to an unmodified mono- nucleosome or higher-order chromatin substrate. **C, D.** Model of BRCA1/BARD1 (BARD1 full-length or RING-IDR) binding with an unmodified mono-nucleosome or higher-order chromatin substrate. **E, F.** Model of BRCA1/BARD1^FL^ binding to ubiquitylated chromatin substrates in a ‘wrapped complex (panel E) or in a complex that spans nucleosome units (panel F). The relative H2A-Ub activity and affinity of the complexes is indicated below each set of panels.

Chemical crosslinking experiments using amine-reactive crosslinkers DSS and BS3 and analyzed by mass spectrometry were carried out to compare complexes formed on unmodified versus H2A K15-Ub-modified nucleosomes **(Supplemental Figure 14)**. To focus on the BARD1 subunit, BRCA1^RING^/BARD1^FL^ was used. There were clear differences in cross-linked products involving BARD1 CTDs, with numerous intermolecular crosslinks (judged by unique crosslinks and peptide spectral mapping counts) to nucleosomes containing H2A K15-Ub but only sparse crosslinks from reactions that included unmodified nucleosomes **(Figure 6E and Supplemental Figure 14C)**. The crosslinks detected are consistent with high-resolution structures of BARD1 CTDs bound to nucleosomes containing H2A K15-Ub and the lack of these in the unmodified nucleosome reaction is consistent with the interaction being critically dependent on the presence of Ub at H2A K15 **(Supplemental Figure 14D)**. There was also increased crosslinking between BARD1 RING and IDR domains to histones in samples containing H2A K15-Ub nucleosomes **(Figure 6E, gray crosslinks and Supplemental Figure 14C, E)**, consistent with the reported higher binding affinity of that complex. Thus, binding of the BARD1 CTDs in response to the presence of H2A K15-Ub facilitates BARD1 RING and IDR interactions that lead to the observed increase in H2A K127-Ub activity.

The BARD1 RING domain and CTDs are separated by ∼300 intrinsically disordered residues, allowing for as much as 1000 Å of separation between the two structured domains. It is therefore possible that the CTDs bind to one nucleosome unit, while the RING domains are recruited to a nearby nucleosome unit on chromatin to facilitate H2A-Ub activity. To test this possibility explicitly, asymmetric di-nucleosome substrates (di-NCPs) where one nucleosome unit contained H2A K15-Ub (H2A^silent^) and an adjacent nucleosome unit contained unmodified H2A with a fluorophore conjugated near its N-terminus to facilitate specific detection in a nucleosome ubiquitylation assay were assembled (H2A^observe^; **Figure 6F and Supplemental Figure 13E-G)**. The asymmetric di-NCP substrate containing H2A K15-Ub as H2A^silent^ was rapidly ubiquitylated by BRCA1^RING^/BARD1^FL^ and this level of activity requires the BARD1 CTDs (H2A^observe^; **Figure 6G and Supplemental Figure 13H)**. Unmodified di-NCPs were ubiquitylated with similar kinetics regardless of the presence or absence of the CTDs. The results confirm that BARD1 CTDs bound to an H2A K15-Ub-containing nucleosome enable RING binding to a neighboring nucleosome and that this configuration leads to high ubiquitylation activity.

Furthermore, deletion of the BARD1 DNA-binding IDR yields a substantial loss of H2A-Ub activity, indicating that all three binding functionalities are required for full activity. Altogether, the results support formation of a higher-order chromatin complex where BARD1 spans adjacent nucleosome units in which at least one contains H2A K15-Ub. The complex is mediated through binding of the BRCA1/BARD1 RING domains and BARD1 CTDs to neighboring nucleosome units and of the BARD1 IDR to intervening linker DNA **(Figure 7F)**.

## Discussion

Structural and biochemical studies have revealed that the two RING domains of heterodimeric BRCA1/BARD1 bind to the histone face of a nucleosome and that this interaction is essential for the ability to modify histone H2A at its C-terminal tail (Hu et al., 2021; Witus, Burrell, et al., 2021). Other studies revealed that BARD1 CTD also binds to the histone face, but this interaction requires Ub-modified H2A, a modification that occurs in response to DNA damage (Becker et al., 2021; Hu et al., 2021; Krais et al., 2021). Herein we have identified an additional region of BARD1 that engages with chromatin, namely the BARD1 IDR proximal to the RING (IDR^Prox^). In concert with the RING domains, BARD1 IDR^Prox^ is sufficient to support highly enhanced H2A-Ub activity in vitro. BARD1 IDR^Prox^ increases the binding affinity of the BRCA1/BARD1 RINGs from a micromolar to nanomolar binding regime, likely critical in cells where BRCA1/BARD1 is in low abundance. Indeed, our cellular data reveal an important role for this region in chromatin recruitment under both basal and DNA-damage conditions and in DNA damage repair. Remarkably, this critical function is carried out by a short stretch of intrinsically disordered BARD1 residues that are highly conserved.

We propose distinct complexes of BRCA1/BARD1 and chromatin substrates that depend on the status of nucleosomal histone PTMs (e.g., unmodified or H2A K15-Ub) **(Figure 7)**. Importantly, while BARD1 CTDs are critical for interactions with nucleosomes that contain H2A K15-Ub, they are dispensable for high-affinity interactions with unmodified nucleosomes **(Figure 7C, D)**.

Additional interactions that enhance H2A-Ub activity are mediated through BARD1 IDR^Prox^ which engages nucleosomal and extra-nucleosomal linker DNA. Our findings indicate that DNA binding plays an auxiliary role in the ability of BRCA1/BARD1 to ubiquitylate H2A by boosting the affinity of the Ub ligase/nucleosome complex. Supporting this, a BARD1 RING-domain mutant (BARD1 P89A/W91A) that disrupts histone binding is unable to support H2A-Ub activity with any chromatin substrate tested despite retaining strong nucleosome binding affinity and intrinsic Ub ligase activity. This indicates that high-affinity binding to nucleosomal DNA by BARD1 IDR^Prox^ is not sufficient for H2A-Ub activity, while the positioning of the RING domains on the histone surface to orient the RING-E2∼Ub complex for site-specific ubiquitin transfer to histone H2A is absolutely required.

BARD1 IDR^Prox^ contains two discrete regions with DNA-binding properties, separated by ∼30 residues. Each DNA-binding region has a high Lys/Arg content while the intervening region contains more negatively charged residues (**Supplemental Figure 3C**). The more C-terminal residues of BARD1 IDR^Prox^ (190-216) appear to be the dominant site for dsDNA engagement while the portion of BARD1 IDR^Prox^ (145-165) directly adjacent to the RING domain also engages dsDNA but shows a pronounced preference for specialized DNA structures such as bubble-DNA that have features found in intermediates in HR (Zhao et al., 2017) and possibly in R-loop structures formed during transcription (Santos-Pereira & Aguilera, 2015). The two regions display different secondary structure propensities, and we note a conserved hydrophobic trio (145-MWF) within the RING-adjacent site that is highly affected by bubble-DNA. Structural information regarding IDR-DNA interactions is sparse, but our findings hint at distinct modes of binding by IDRs rather than merely electrostatic, non-specific associations. To that point, BARD1 IDR^Prox^ appears to possess small regions with specialized binding modes that are linked to BRCA1/BARD1 function.

Preferential binding to bubble-DNA serves to inhibit H2A-Ub activity in our assays where the specialized DNA species was present in *trans*. This suggests the intriguing possibility that proximity or high local concentrations of such species might modulate the modification of H2A by BRCA1/BARD1 in cells. Furthermore, binding of BARD1 IDR^Prox^ to nucleosomal DNA or to specialized nucleic acid structures could serve to control its spatial and temporal involvement in HR and other processes where such specialized DNA structures may be present (e.g., stalled replication fork protection, R-loop resolution) (Hatchi et al., 2015; Schlacher et al., 2012).

Additionally, BRCA1/BARD1 is present in higher-order protein assemblies that are known to bind to chromatin via distinct nucleosome interfaces that may be formed by other proteins in the complex (Belotserkovskaya et al., 2020; Bochar et al., 2000; Huen et al., 2010; Savage & Harkin, 2015). Understanding the combined binding contributions of BRCA1/BARD1 and other proteins in such complexes to nucleosome association and H2A-Ub activity is an important future goal.

Data presented here provide direct experimental evidence in support of an extended BRCA1/BARD1/chromatin complex that spans nucleosomal units. Such a complex is enabled by the presence of H2A K15-Ub, an early mark of DNA damage, which engages the BARD1 CTDs while the enzymatic RING domains occupy an adjacent/nearby nucleosome unit **(Figure 7E, F)**. The 300-residue BARD1 IDR allows extensive separation between the enzymatic RING domains and BARD1 CTDs while itself engaging in high-affinity interactions with nucleosomal/linker DNA. Such a configuration may allow the RING domains to reach nucleosome units even further away than an adjacent unit while anchored to chromatin via the BARD1 CTDs. This situation could lead to BRCA1/BARD1-dependent H2A-Ub of multiple nucleosomes in the vicinity of a H2A K15-Ub anchor nucleosome. Additionally, the ability to reach across nucleosome units in chromatin provides a platform to scaffold higher-order complexes across long molecular distances.

As H2A K15-Ub is specifically a mark of DNA damage, this model will not be in play for other BRCA1/BARD1-dependent processes on chromatin such as its transcriptional repression and activation activities. Results presented here show that the DNA-binding BARD1 IDR^Prox^ is required for enhanced H2A-Ub activity on unmodified nucleosomes. An important question going forward is what other histone PTMs influence the chromatin binding and H2A-Ub activity of BRCA1/BARD1. We previously reported that methylation at H3 K79, a mark associated with actively transcribed chromatin, inhibits BRCA1/BARD1-mediated H2A-Ub activity (Witus, Burrell, et al., 2021). This may be attributable to the location of H3 K79 near the BARD1-histone interface. A comprehensive understanding of histone PTMs that influence BRCA1/BARD1 nucleosome binding, higher-order chromatin complex formation, and H2A-Ub activity will provide valuable insight into its functions and regulation in different biological processes.

Mostly missing from our discussion is the huge BRCA1 subunit. Its only established direct point-of-contact to chromatin is via its N-terminal RING domain, although at least one section of its enormous IDR has DNA binding activity in vitro (Mark et al., 2005). BRCA1/BARD1 serves as a large protein interaction hub that forms numerous distinct complexes, many of which involve BRCA1 as the binding partner. In our in vitro investigation, the entirety of BRCA1 apart from its enzymatic RING domain, was dispensable for H2A-Ub activity. That said, we note that full-length BRCA1/BARD1 displayed lower H2A-Ub activity than complexes containing truncated BRCA1 (Δ11q or RING) and offer two hypotheses for the observation. First, BRCA1^FL^ but neither truncation contains a DNA-binding region that supports lower H2A-Ub activity than BARD1 IDR^Prox^. The presence of strong DNA-binding regions in both BRCA1 and BARD1 in the full-length complex could set up a competition between these regions for binding to nucleosomal DNA, leading to lower apparent H2A-Ub activity. Alternatively, or in addition, possible inhibitory interactions between BRCA1 and BARD1 subunits might have been alleviated upon BRCA1 truncation. Although such effects could be artifactual, we suggest that the status of the BRCA1 subunit in terms of its binding partners could provide an additional layer of modulation of BRCA1/BARD1 H2A-Ub activity. Occupancy of regions of BRCA1 and BARD1 by unique binding partners may occlude certain nucleosome-binding regions, resulting in inhibition or enhancement of chromatin binding and H2A-Ub activity. Further investigation into chromatin complexes containing full-length BRCA1/BARD1 and the many protein assemblies and binding partners that are required to execute its cellular functions is warranted.

In conclusion, our results establish an extensive multivalent network of interactions that facilitate BRCA1/BARD1 chromatin association and H2A-Ub activity. These interactions are likely critical for many functions of BRCA1/BARD1 in the nucleus that are not limited to its H2A-Ub activity and may be disrupted by mutations that cause cancer.

## Acknowledgements

We thank K. Luger (University of Colorado Boulder) for sharing plasmids for the 147-bp and NLE-trimer ‘601’ DNA, S. Tan (Penn State University) for sharing the plasmid for the 185-bp ‘601’ DNA, and G. Debelouchina (UC San Diego) for the MMTV DNA plasmid. We thank D. Veelser for providing access to equipment, and M. Morgan for insights into dichloroacetone crosslinking of H2A-Ub. S.R.W., L.M.T., K.E.K., P.S.B., and R.E.K. were supported by the NIH (R01CA260834). R.E.K. is the Edmond H. Fischer/Washington Research Foundation Endowed Chair in Biochemistry. This work and A.Z. was supported in part by the University of Washington’s Proteomics Resource (UWPR95794). A.Z. and T.N.D. were also supported by the NIH (P41GM103533). W.Z. was supported by a Young Investigator Award from Max and Minnie Tomerlin Voelcker Fund, the Cancer Prevention and Research Institute of Texas (RP210102), and the NIH (R01GM141091). D.B.W. was supported by the NIH (R00-HD090201).

## Author Contributions

S.R.W., W.Z., P.S.B., and R.E.K. conceived of the study. S.R.W., W.L., L.M.T., M.W., W.Z., and K.E.K. performed experiments and analyzed data. D.B.W. analyzed data. A.Z. performed chemical crosslinking and mass spectrometry and associated data processing under the supervision of T.N.D. The manuscript was written by S.R.W., P.S.B., and R.E.K., with input from all co-authors.

## Supplemental Figures

**Supplemental Figure 1.**
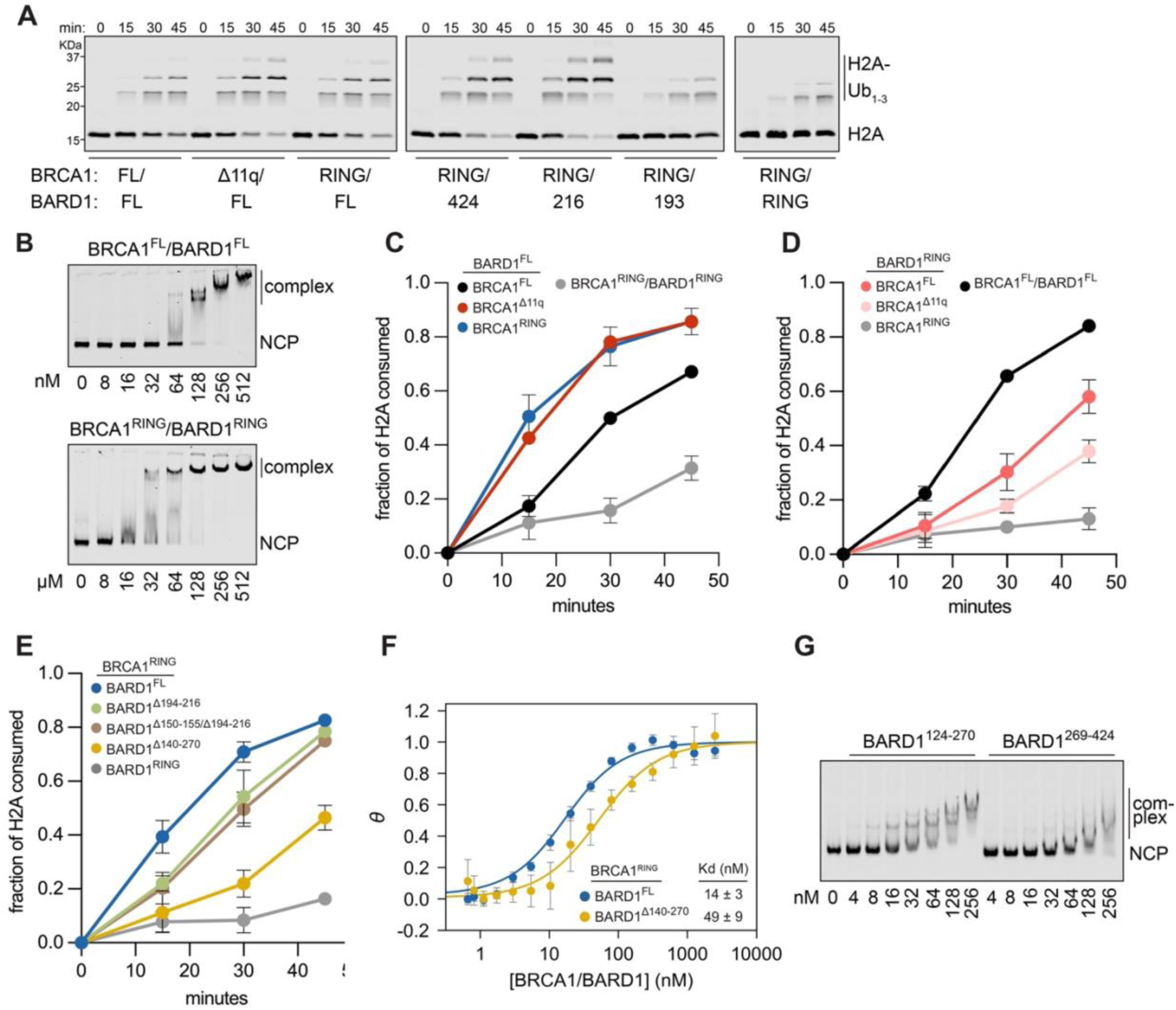
Characterization of BRCA1/BARD1 constructs. **A.** Representative time-course nucleosome ubiquitylation assays monitoring H2A-Ub efficiency of different BRCA1/BARD1 constructs quantified in Panel E and Figure 1B. These data are representative of the basic assay set-up for other time-course H2A-Ub assays in this study. **B.** Electrophoretic mobility shift assay (EMSA) comparing the nucleosome binding of full-length BRCA1/BARD1 (top) to the minimal RING heterodimer (bottom). Data shown are representative of n=2 independent experiments. **C-E.** Quantification of H2A-Ub assays using the indicated BRCA1/BARD1 constructs. Data show the mean; error bars are ± 1-s.d. of n=3 independent experiments. **F.** Comparison of nucleosome binding affinity from fluorescence-quenching based measurements using the indicated BRCA1/BARD1 constructs. Data show the mean; error bars are ± 1-s.d. of n=4 independent experiments. **G.** EMSA monitoring nucleosome binding by the indicated BARD1 IDR fragments. Data shown are representative of n=2 independent experiments.

**Supplemental Figure 2.**
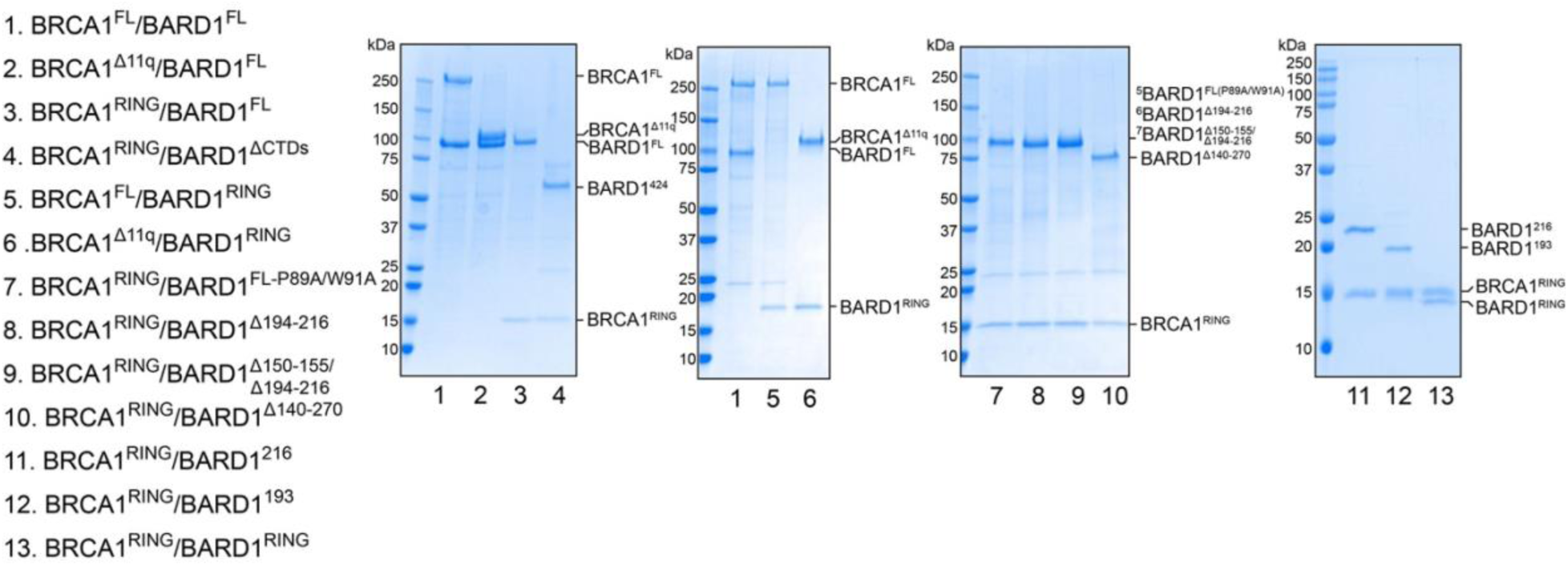
**Coomassie-stained SDS-PAGE gels of BRCA1/BARD1 constructs used in this study.**

**Supplemental Figure 3.**
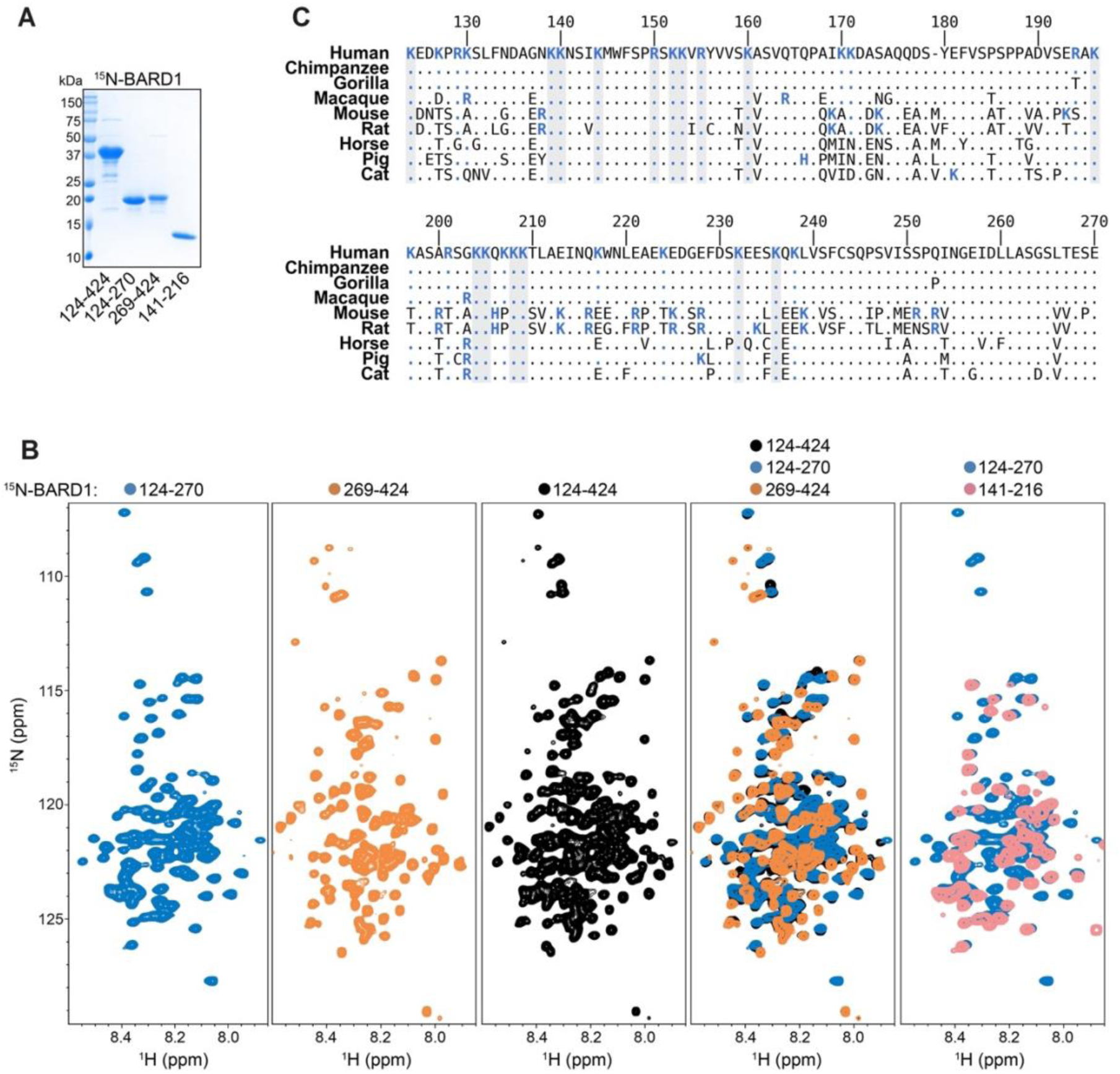
Characterization of the BARD1 intrinsically disordered region. **A.** Coomassie-stained SDS-PAGE gel of BARD1 fragments used in NMR experiments. **B.** ^1^H^15^N-HSQC NMR spectra of the indicated ^15^N-labelled BARD1 fragments. **C.** Multiple-sequence alignment of BARD1 124-270 with a representative subset of homologs. Residue identities that are conserved to the human sequence are indicated by the presence of a period in place of a residue letter. Basic residues (Lys, Arg, His) are color coded in blue, and basic positions that are fully conserved across all species are highlighted with a gray background.

**Supplemental Figure 4.**
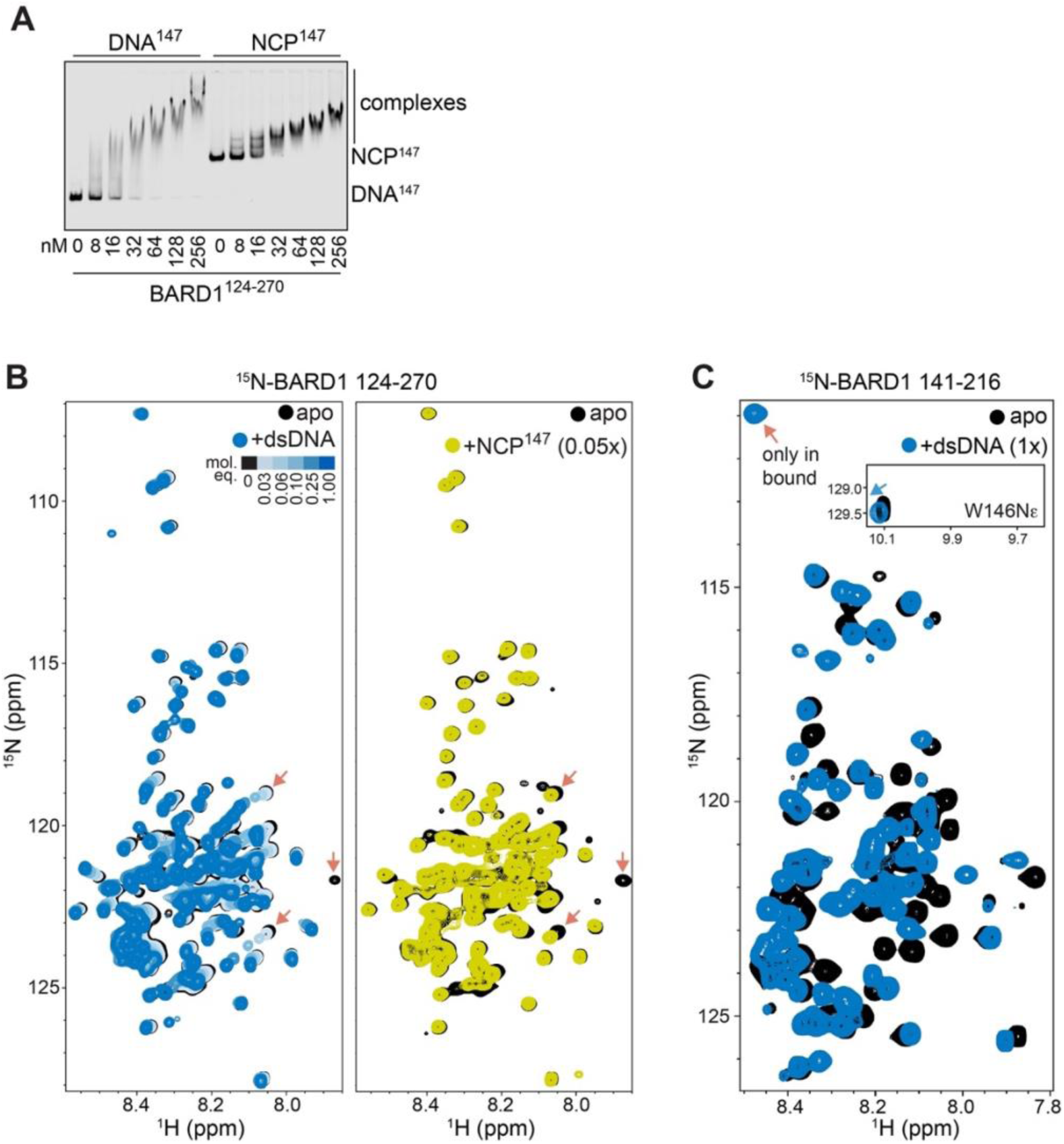
NMR analysis of BARD1 IDR-DNA and IDR-nucleosome complexes. **A.** EMSA comparing DNA and nucleosome binding by BARD1 124-270. Data shown are representative of n=2 independent experiments. **B.** ^1^H^15^N-HSQC NMR spectra of ^15^N-BARD1 124-270, titrating a 36-bp dsDNA fragment (left), and a single titration point with nucleosomes (right). The red arrows point to signals that are similarly affected between the dsDNA- and nucleosome-bound samples. **C.** ^1^H^15^N-HSQC NMR spectra of ^15^N-BARD1 141-216 with a 36-mer dsDNA fragment. Data from panel C are also shown in Supplemental Figure 10A.

**Supplemental Figure 5.**
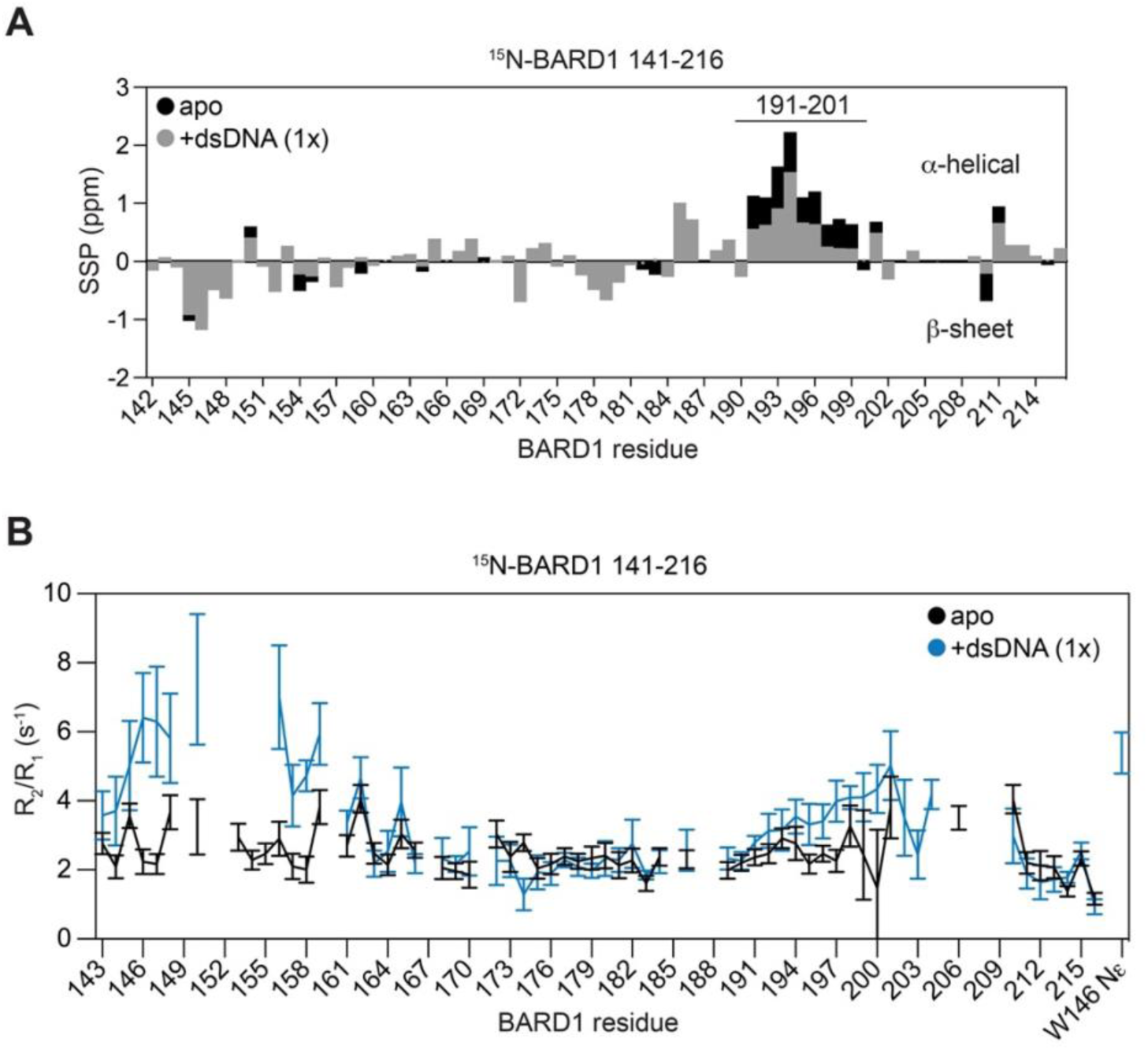
Changes in dynamics and secondary structure of the BARD1 IDR bound to DNA. **A.** Changes in secondary structural propensity (SSP) of ^15^N-BARD1 141-216 when bound to a 36-bp dsDNA fragment. The black bars for apo BARD1 are behind the gray bars where DNA was added. **B.** T1/T2 dynamics measurements for amide signals of ^15^N-BARD1 141-216 in apo form and bound to dsDNA (1:1 molar equivalent complex). Data from panel B are also shown in Supplemental Figure 10C.

**Supplemental Figure 6.**
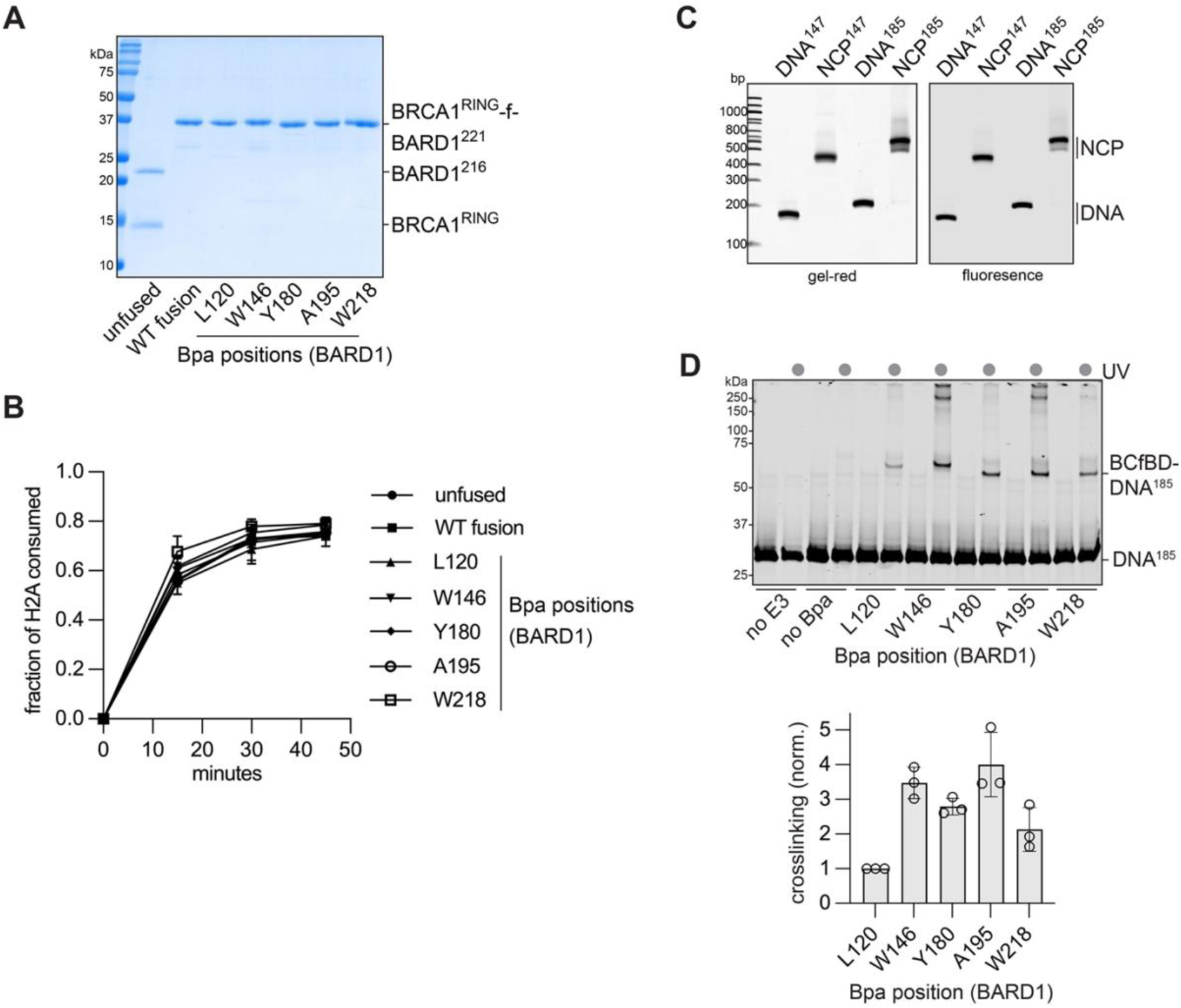
UV-induced Bpa crosslinking of BRCA1/BARD1 to nucleosomal DNA. **A.** Coomassie-stained SDS-PAGE gel of unfused BRCA1^RING^/BARD1^216^ heterodimers and BRCA1-f-BARD1^221^ constructs with and without Bpa incorporation. **B.** H2A-Ub activity of unfused heterodimers compared to wild-type and Bpa-incorporated BRCA1-f-BARD1^221^. Data show the mean; error bars are ± 1-s.d. of n=3 independent experiments. **C.** Native gel of fluorescently labelled DNA and nucleosomes used in Bpa crosslinking experiments. **D.** Representative data of in-gel fluorescence signal from of a UV-induced crosslinking reaction to free 185-bp ‘601’ DNA (DNA^185^, top) and quantification of crosslinking experiments (bottom). The intensity.y of each crosslinked band was normalized to the L120Bpa crosslinked band intensity for each replicate experiment. Data bars show the mean; error bars are ± 1-s.d. and the open circles are the values of individual replicates for of n=3 independent experiments.

**Supplemental Figure 7.**
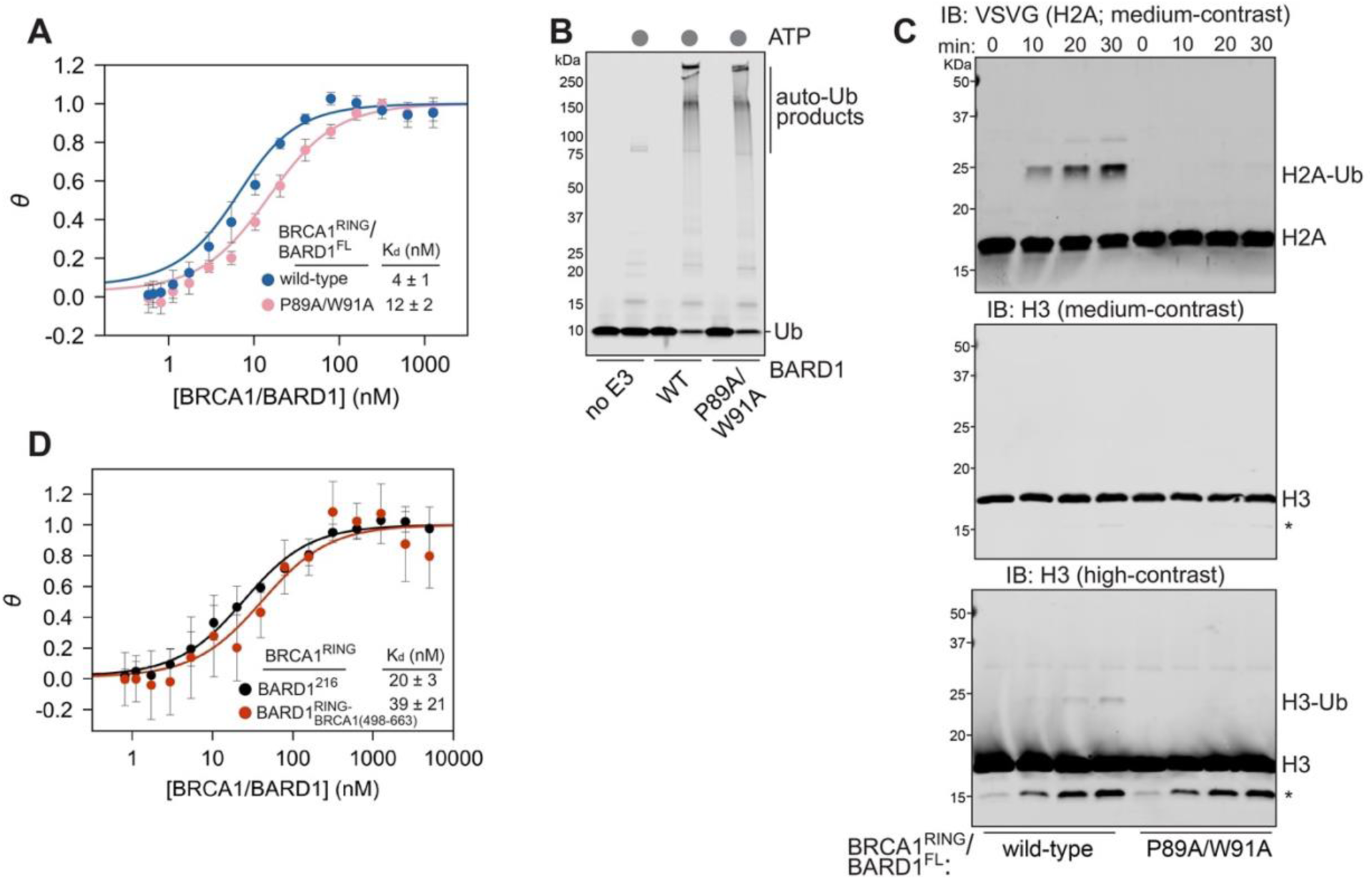
Nucleosome binding and Ub ligase activity of BARD1 RING mutants and DNA-binding chimeric constructs. **A.** Comparison of nucleosome binding affinity using the indicated BRCA1/BARD1 constructs. Data show the mean; error bars are ± 1-s.d. of n=3 independent experiments. **B.** Auto-ubiquitylation assay using BRCA1^RING^/BARD1^FL^ (wild-type or P89A/W91A). The assay monitors depletion of Ub and appearance of high molecular weight auto-Ub products in the presence of BRCA1/BARD1. Data shown is representative of n=2 independent experiments. **C.** Nucleosome ubiquitylation assay using the indicated BRCA1/BARD1 constructs and blotting for VSVG-tagged H2A (top) and H3 (middle and bottom). The asterisk denotes a proteolysis product of H3 that appears over the time-course. **D.** Comparison of nucleosome binding affinity using the indicated BRCA1/BARD1 constructs. Data show the mean; error bars are ± 1-s.d. of n=2 independent experiments.

**Supplemental Figure 8.**
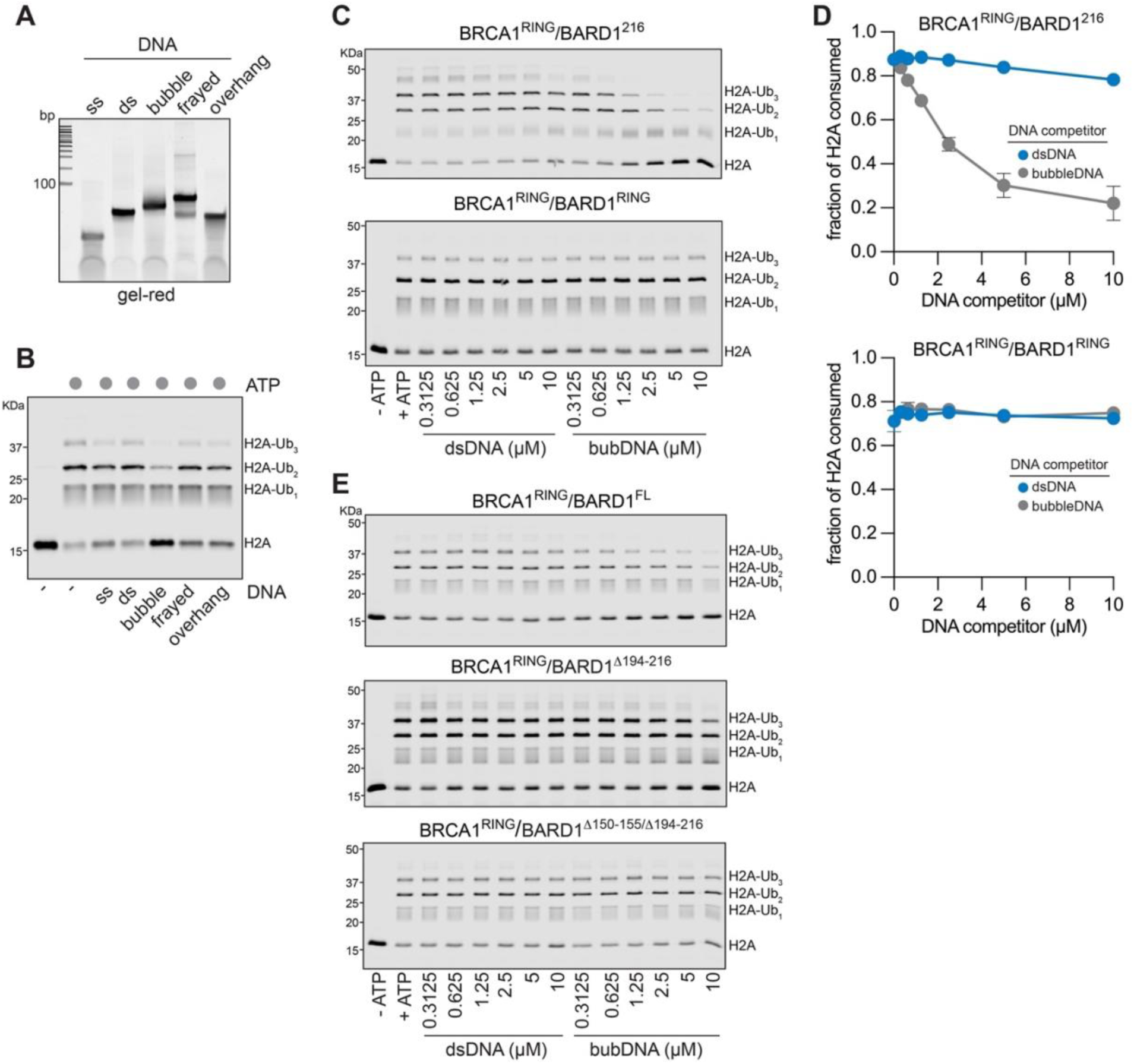
Assembly of competitor DNA fragments and inhibition of H2A-Ub activity. **A.** DNA-stained native-gel of annealed DNA competitor fragments. Amounts loaded were normalized to show similar band-intensity for each fragment. **B.** Representative gel data of H2A-Ub inhibition assay using full-length BRCA1/BARD1 (100 nM) and the indicated DNA competitor fragment (2.5 mM) **C.** Representative gel data of single time-point (20 min) H2A-Ub inhibition assays using BRCA1^RING^/BARD1^216^ (top, 50 nM E3) and BRCA1^RING^/BARD1^140^ (bottom, 750 nM E3) and indicated amounts of dsDNA and bubble-DNA competitors. **D.** Quantification of H2A-Ub DNA-competitor inhibition assays from panel b using the indicated BRCA1/BARD1 constructs. Data show the mean; error bars are ± 1-s.d. of n=3 independent experiments. **E.** Representative gel data of single time-point H2A-Ub inhibition assays using BRCA1^RING^/BARD1^FL^ (top, 50 nM E3, 12 min endpoint), BRCA1^RING^/BARD1^Δ194-216^ (middle, 50 nM E3, 20 min endpoint), and BRCA1^RING^/BARD1^Δ150-155/Δ194-216^ (bottom, 50 nM E3, 20 min endpoint) and indicated amounts of dsDNA and bubble-DNA competitors.

**Supplemental Figure 9.**
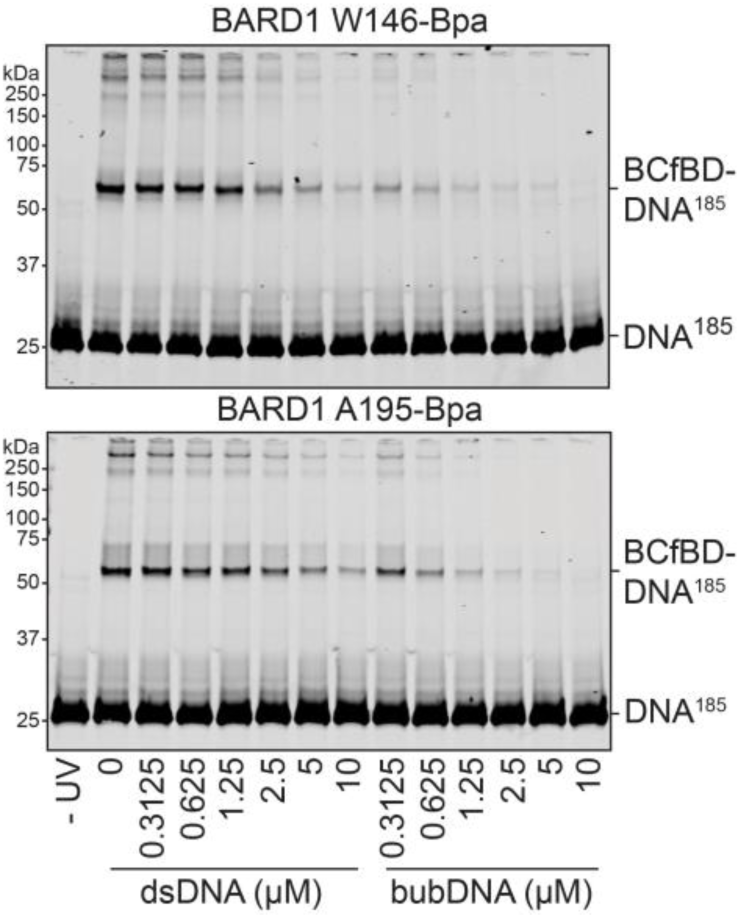
Inhibition of UV-induced Bpa crosslinking of BRCA1/BARD1 to nucleosomal DNA. Representative data of in-gel fluorescence signal from a UV-induced Bpa crosslinking reaction between W146Bpa (top) and A195Bpa (bottom) BRCA1-f-BARD1^221^ to 185-bp ‘601’ nucleosomes (NCP^185^) in the presence of dsDNA and bubble-DNA competitors. Quantified data from n=3 replicate experiments are shown in Figure 4C.

**Supplemental Figure 10.**
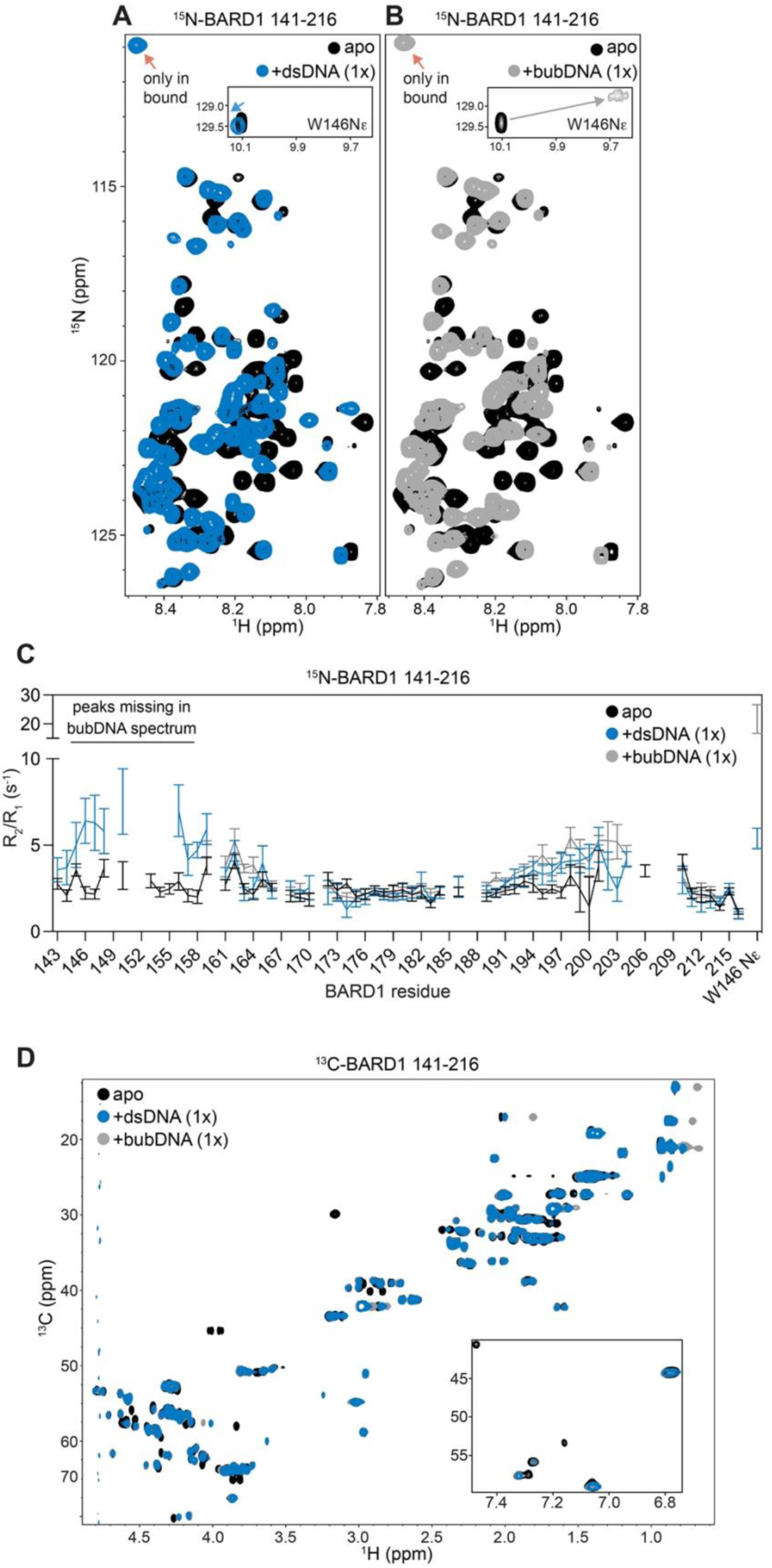
Comparison of dsDNA and bubble-DNA binding to the BARD1 IDR by NMR. **A, B.** ^1^H^15^N-HSQC NMR spectra of ^15^N-BARD1 141-216 with a 36-mer dsDNA fragment (A) or a 36-mer bubble-DNA fragment with 8-bp of mismatch (B). Both complexes were assembled at 1:1 molar equivalent. The W146Nε signals are inlaid, and the red arrows point to a signal that is present exclusively in the DNA-bound spectra. Data from panel A are also shown in Supplemental Figure 4C. **C.** T1/T2 dynamics measurements for amide signals of ^15^N-BARD1 141-216 in apo form and bound to dsDNA or bubble-DNA (1:1 molar equivalent complex). A subset of data from panel C (apo and +dsDNA) are also shown in Supplemental Figure 5B. **D.** ^1^H^13^C-HSQC NMR spectra of ^13^C-BARD1 141-216 with a 36-mer dsDNA fragment (blue), or a 36-mer bubble-DNA fragment with 8-bp of mismatch (gray). Both complexes were assembled at 1:1 molar equivalent.

**Supplemental Figure 11.**
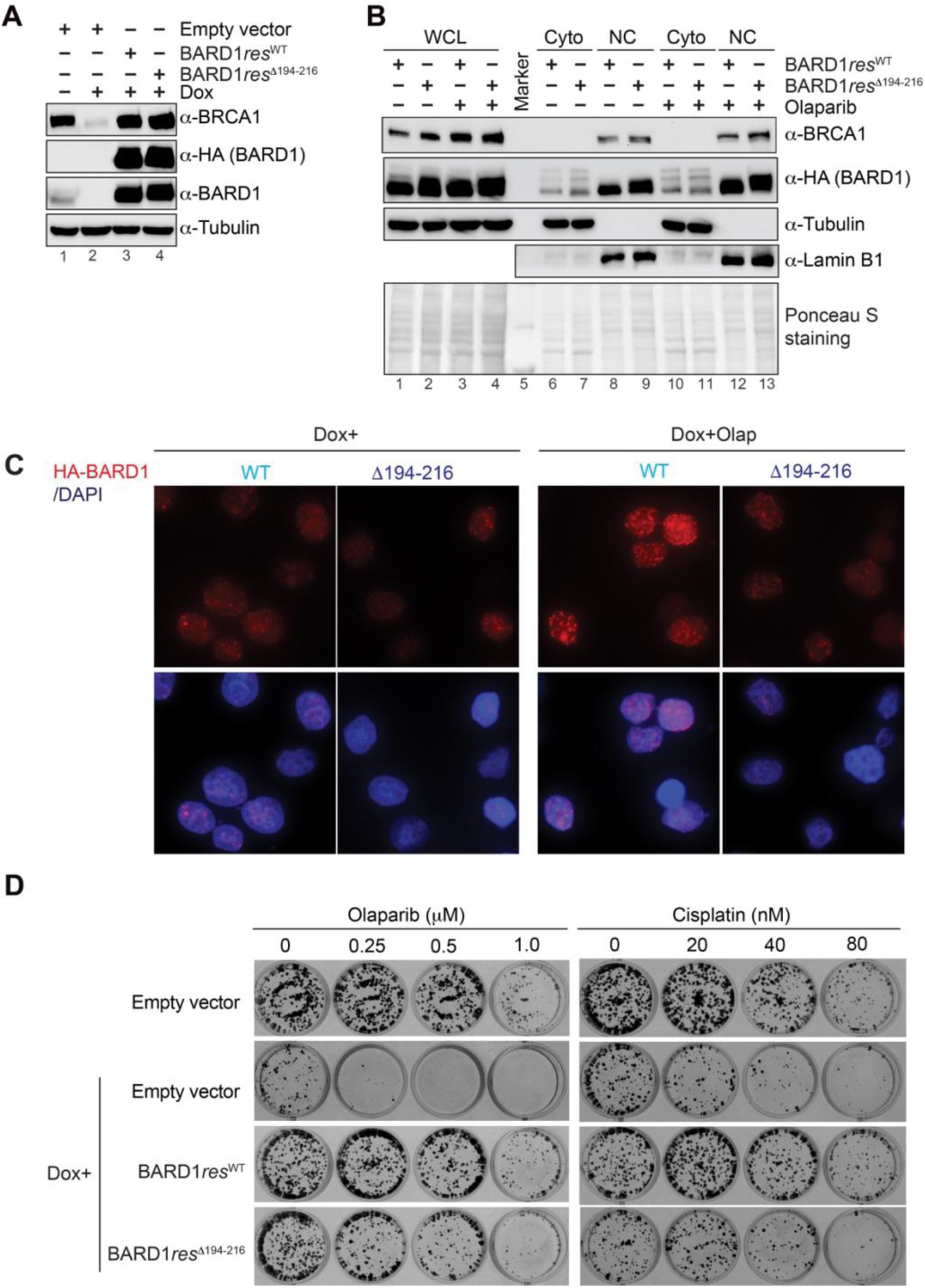
Cellular studies of BARD1 DNA-binding mutant. **A.** Western blot analysis to detect BRCA1, BARD1 and HA-BARD1 of the whole-cell lysate from HeLa-shBARD1 cells stably expressing wild type or Δ194-216 mutant of HA-BARD1, where endogenous BARD1 was depleted by doxycycline induced shBARD1 expression. The tubulin blot was included as a loading control. **B.** Western blot analysis to detect BRCA1 and HA-BARD1 in the whole-cell lysate (WCL), cytoplasmic fraction (Cyto) and nuclear extraction (NC) from HeLa-shBARD1 cells stably expressing wild type or Δ194-216 mutant of HA-BARD1, where endogenous BARD1 was depleted by doxycycline treatment. The further fractionations of the nuclei isolated are shown in Figure 5A. Tubulin and Lamin B1 were included as loading controls and as indication of the Cyto and NC fractions, respectively. **C.** Representative micrographs of HA-BARD1 foci (red) in the nucleus of HeLa-shBARD1 cells stably expressing wild type or Δ194-216 mutant of HA-BARD1 with or without the treatment of 10 mM olaparib 24h (left panels). Blue: DAPI. Cells with >5 HA-BARD1 foci and number of HA- BARD1 foci per nuclear were quantified (reported in Figure 5B). **D.** Representative images for cell survival data quantified in Figure 5C.

**Supplemental Figure 12.**
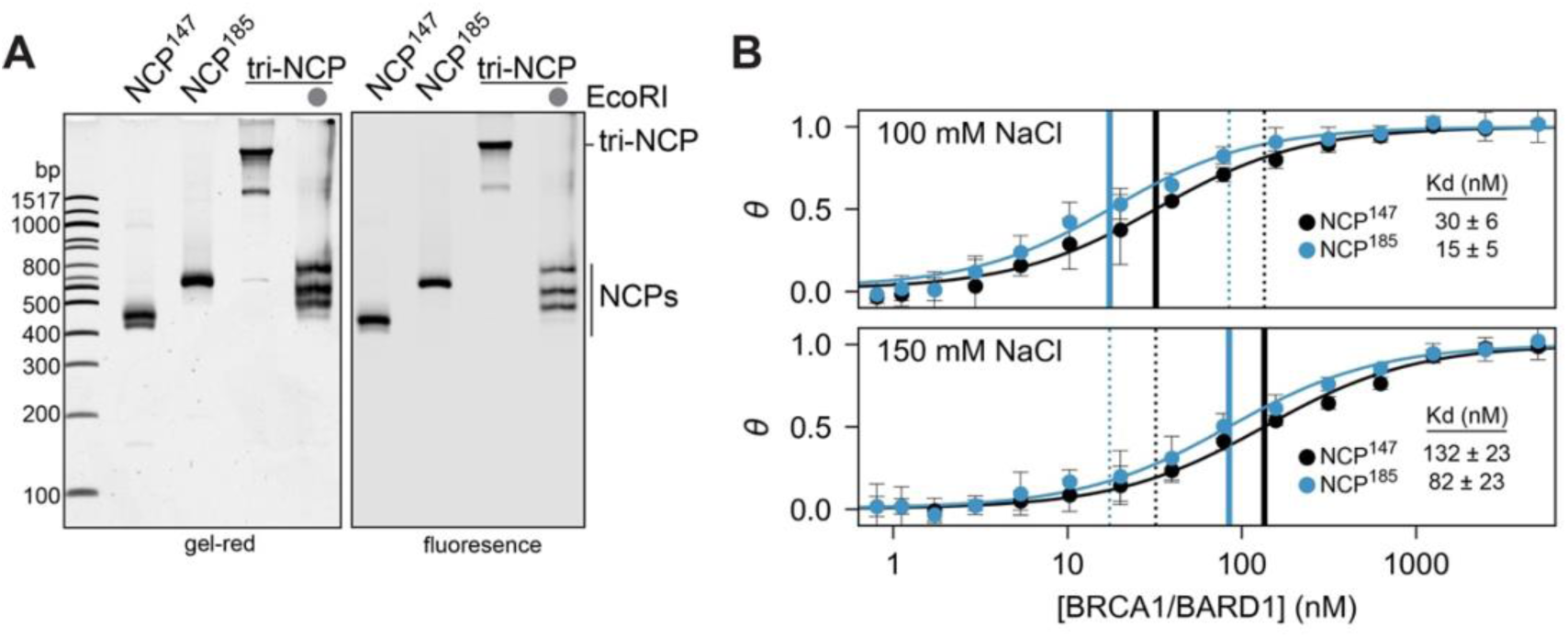
Analysis of chromatin substrates. **A.** DNA-stained (left) and fluorescence (right) native gel of chromatin substrates used for H2A- Ub activity assays in Figure 5b and c. EcoRI digestion of tri-NCPs is used to confirm octamer binding saturation. **B.** Comparison of nucleosome binding affinity between BRCA1^RING^/BARD1^216^ and NCP^147^ and NCP^185^ substrates at 100 mM (top) and 150 mM (bottom) buffer-salt concentration (NaCl). The solid vertical lines denote the position of K_d_ values for the indicated buffer-salt concentration, and the dotted vertical lines denote the position of K_d_ values from the other buffer-salt concentration. Data show the mean; error bars are ± 1-s.d. of n=3 independent experiments using both NCP^147^ and NCP^185^, with an additional replicate (total of n=4) for NCP^147^.

**Supplemental Figure 13.**
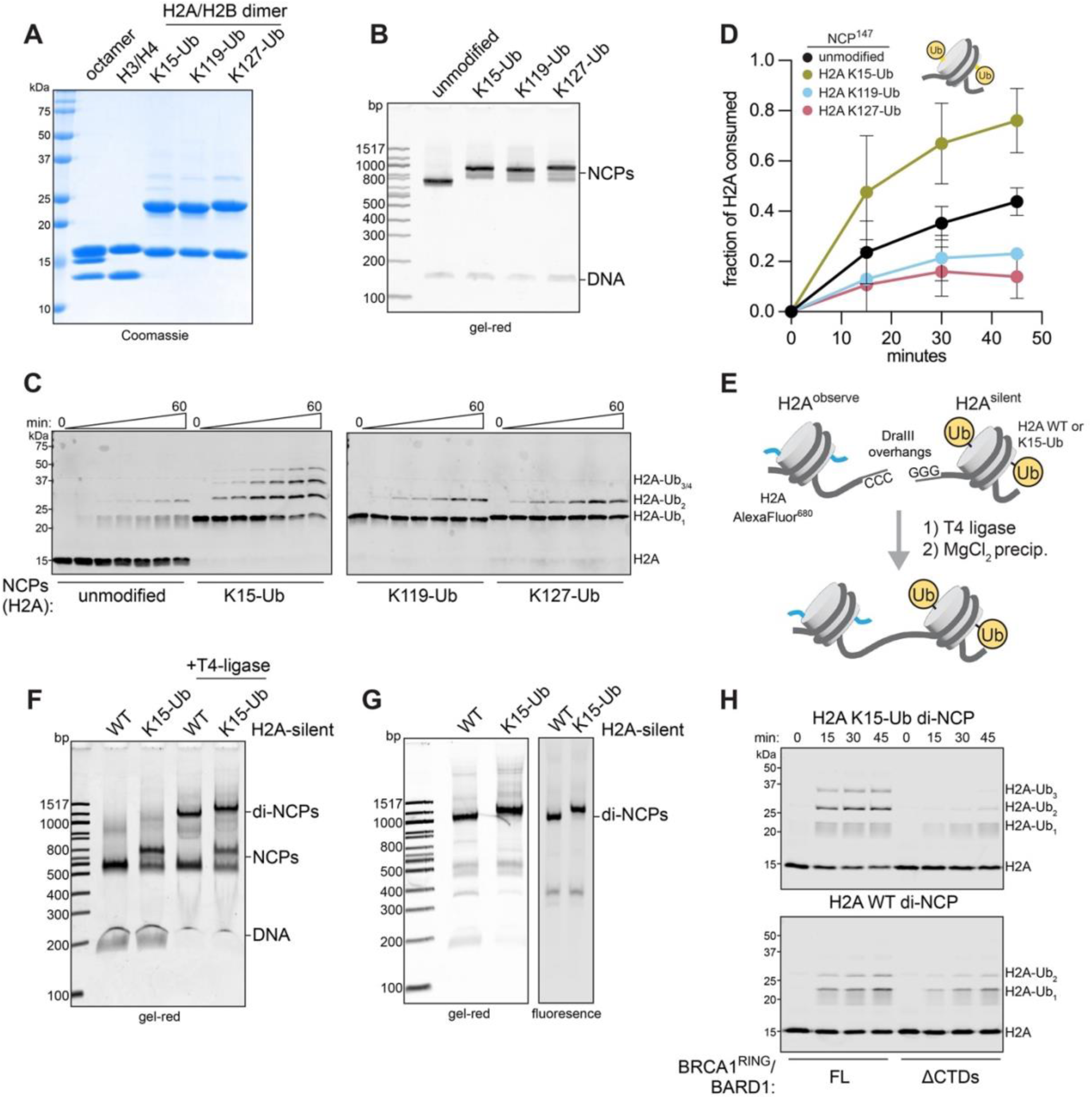
Design, assembly, and quality control of H2A K15-Ub chromatin substrates. **A.** Coomassie-stained SDS-PAGE gel of wild-type histones and dichloroacetone linked H2A- Ub/H2B dimers. **B.** DNA-stained native gel of mono-nucleosomes with and without H2A-Ub incorporation at the indicated positions. **C.** Example data of H2A-Ub activity assay using full-length BRCA1/BARD1 and preinstalled H2A-Ub NCP substrates. Western blots were probed for H2A. **D.** Quantification of time-course H2A-Ub assays using unmodified or pre-installed H2A-Ub chromatin substrates and BRCA1^FL^/BARD1^FL^. **E.** Schematic of asymmetric di-NCP assembly strategy. **F.** DNA-stained native gel of di-NCP ligation reaction. **G.** DNA-stained (left) and fluorescence (right) native gels of di-NCP substrates after purification by MgCl_2_ precipitation. **H.** Example gel data of H2A-Ub activity assay using di-NCP substrates and indicated BRCA1/BARD1 constructs.

**Supplemental Figure 14.**
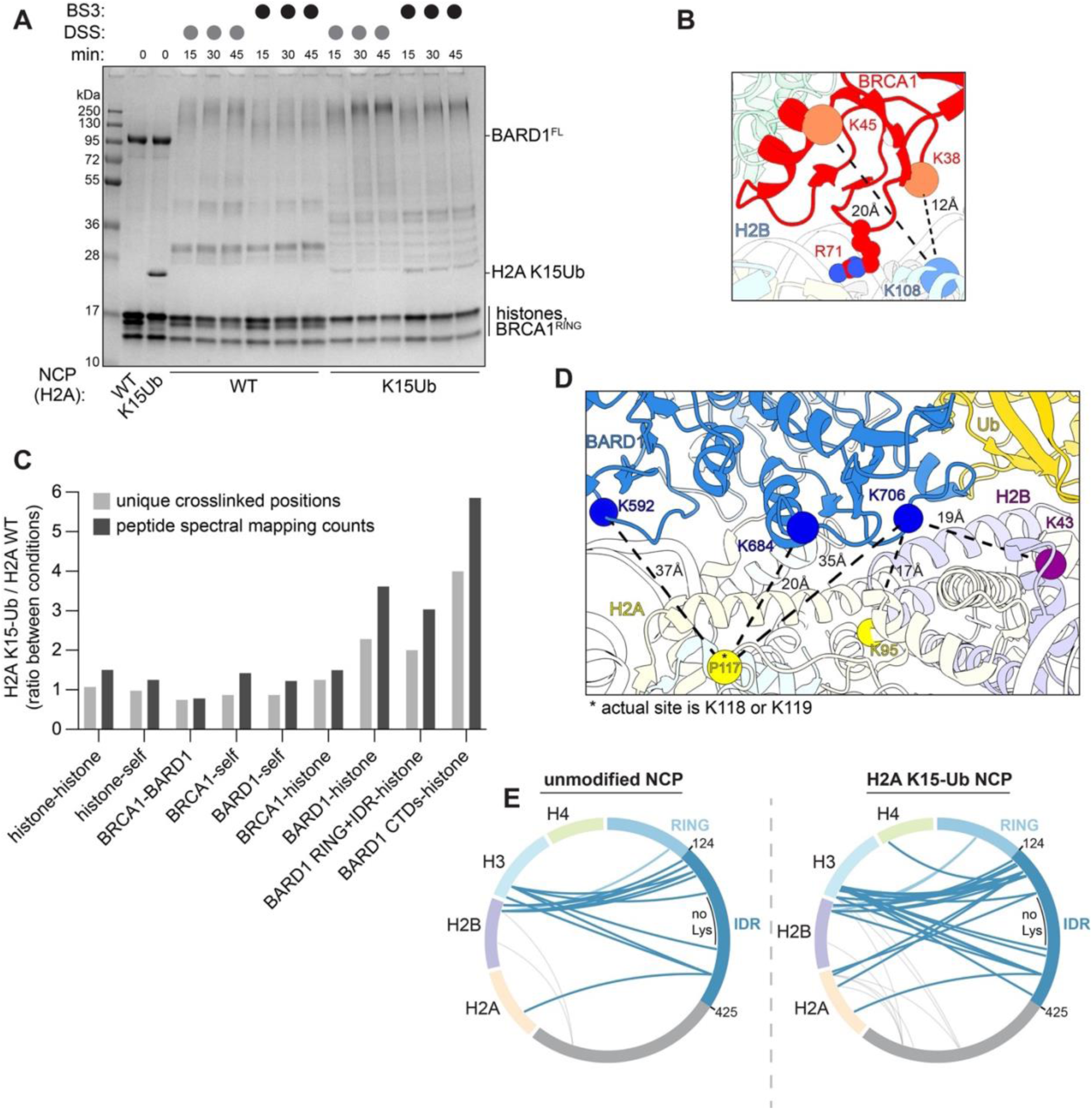
Analysis of BRCA1/BARD1 chromatin complexes by chemical crosslinking and mass-spectrometry. **A.** Coomassie-stained SDS-PAGE gel of chemical crosslinking reactions used in mass spectrometry analysis **B.** Crosslinks observed from the RING domain of BRCA1 to ordered regions histones mapped on a high-resolution structure of the complex (PDB: 7JZV). Cα-Cα distances are indicated in the figure. **C.** Quantitative comparison of crosslinking from reactions with unmodified and H2A K15-Ub NCPs. Different subsets of crosslinks were quantified for the number of unique observed crosslinks and total peptide spectral mapping counts for all crosslinks within a given subset. **D.** Crosslinks observed from the CTDs of BARD1 to ordered regions histones in the H2A K15- Ub NCP sample mapped on a high-resolution structure (PDB: 7E8I). Cα-Cα distances are indicated. The P117 Cα serves as a proxy for the true crosslinking sites of K118 and K119 that are not modelled, denoted by an asterisk. **E.** Intermolecular crosslinks observed by chemical crosslinking and MS analysis between BARD1 and histones using wild-type (left) and H2A K15-Ub (right, separated by dashed line) 147-bp ‘601’ nucleosomes and BRCA1^RING^/BARD1^FL^ heterodimers. Crosslinks to histones emanating from the BARD1 RING-IDR region are highlighted in blue. A lysine-depleted region of the BARD1 IDR is labelled and indicated by a black bar.

## Tables

**Extended Data Table I.** Details about BRCA1/BARD1 constructs used in this study.

| Construct | BRCA1 bounds* | BRCA1 tags | BARD1 bounds* | BARD1 tags | Expression system |
| --- | --- | --- | --- | --- | --- |
| BRCA1<sup>FL</sup>/<br>BARD1<sup>FL</sup> | 1-1863 | N-FLAG | 1-777 | N-TwinStrepII | *Sf9* |
| BRCA1<sup>Δ11q</sup>/<br>BARD1<sup>FL</sup> | 1-264,<br>1366-1863 | N-FLAG | 1-777 | N-TwinStrepII | *Sf9* |
| BRCA1<sup>RING</sup>/<br>BARD1<sup>FL</sup> | 1-112 | N-FLAG | 1-777 | N-TwinStrepII | *Sf9* |
| BRCA1<sup>RING</sup>/<br>BARD1<sup>ΔCTDs</sup> | 1-112 | N-FLAG | 1-424 | N-TwinStrepII | *Sf9* |
| BRCA1<sup>FL</sup>/<br>BARD1<sup>RING</sup> | 1-1863 | N-FLAG | 1-140 | N-TwinStrepII | *Sf9* |
| BRCA1<sup>Δ11q</sup>/<br>BARD1<sup>RING</sup> | 1-264,<br>1366-1863 | N-FLAG | 1-140 | N-TwinStrepII | *Sf9* |
| BRCA1<sup>RING</sup>/<br>BARD1<sup>Δ194-216</sup> | 1-112 | N-FLAG | 1-193,<br>217-777 | N-TwinStrepII | *Sf9* |
| BRCA1<sup>RING</sup>/<br>BARD1<sup>Δ150-155/Δ194-216</sup> | 1-112 | N-FLAG | 1-149,<br>156-193,<br>217-777 | N-TwinStrepII | *Sf9* |
| BRCA1<sup>RING</sup>/<br>BARD1<sup>Δ140-270</sup> | 1-112 | N-FLAG | 1-139,<br>271-777 | N-TwinStrepII | *Sf9* |
| BRCA1<sup>RING</sup>/<br>BARD1<sup>216</sup> | 1-110 | N-6xHis | 26-216 | none | *E. coli* |
| BRCA1<sup>RING</sup>/<br>BARD1<sup>193</sup> | 1-110 | N-6xHis | 26-193 | none | *E. coli* |
| BRCA1<sup>RING</sup>/<br>BARD1<sup>RING</sup> | 1-110 | N-6xHis | 26-140 | none | *E. coli* |

\* BRCA1 UniProt sequence ID: P38398-1; BARD1 UniProt sequence ID: Q99728-1

## Methods

### Protein, nucleosome, and experimental reagent production

*Ubiquitylation machinery:* Human E1 (UBA1), E2 (UBE2D3), and ubiquitin (wild-type and Q2C mutant) were expressed and purified as previously described(Brzovic et al., 2003; Christensen et al., 2007; Lazar et al., 1997). For a full list of BRCA1/BARD1 truncation constructs, their residue bounds, purification/epitope tags, and expression system, see Extended Data Table I.

*BRCA1/BARD1 from E. coli:* For purification of truncated BRCA1/BARD1 heterodimers from *E. coli* expression (BRCA1^RING^/BARD1^RING^, BRCA1^RING^/BARD1^193^, and BRCA1^RING^/BARD1^216^), plasmids containing 6xHis-tagged BRCA1 (pCOT7n) and untagged BARD1 (pET28n) were co-transformed into *E. coli* BL21 (DE3) cells, grown in LB media at 37 °C to OD_600nm_ of 0.6-0.8, supplemented with 100 μM ZnCl_2_ and induced with 0.2 mM IPTG for ∼16 hours at 16 °C. The cell pellets were resuspended in 30 mL of Ni^2+^ start buffer (25 mM Tris-HCl pH 7.5, 500 mM NaCl, 10 mM imidazole) supplemented with DNase, RNase, EDTA-free protease inhibitors (Roche) and 1 mM PMSF, lysed by French press, centrifuged, and the supernatant was applied to a 5 mL HisTrap FF crude column (Cytiva). After extensive washing with Ni^2+^ start buffer containing 10 mM imidazole and 30 mM imidazole, protein was eluted using the same buffer supplemented with 500 mM imidazole. The BRCA1^RING^/BARD1^RING^ and BRCA1^RING^/BARD1^193^ constructs were concentrated and further purified using a 120 mL Superdex 75 column on an AKTA FPLC system in SEC buffer (25 mM HEPES-NaOH pH 7.5, 150 mM NaCl, 1 mM DTT; GE Healthcare). For the BRCA1^RING^/BARD1^216^, HisTrap elutions were dialyzed into ion-exchange buffer (25 mM Tris-HCl pH 7.5, 250 mM NaCl, 1 mM DTT) and applied to a 5 mL HiTrap SP HP column (Cytiva) and eluted over a 12 column-volume (CV) gradient (0.25-1 M NaCl). Peak fractions were concentrated, aliquoted, and flash frozen.

For Bpa crosslinking experiments, 6xHis-BRCA1^1-104^-(GS)_6_-BARD1^26-221^ was sub-cloned using Gibson assembly into a pET28n vector. Constructs for Bpa incorporation were generated via site-directed mutagenesis to install amber stop codons in place of BARD1 L120, W146, Y180, A195, and W218. Bpa-mutant Pet28n-6xHis-BRCA1^1-104^-(GS)_6_-BARD1^26-221^ plasmids were co-transformed with a pEVOL-pBpF plasmid encoding an Bpa aminoacyl tRNA synthetase (gift from P. Schultz, Addgene #31190) in *E. coli* BL21 (DE3) cells(Chin Jason W. et al., 2002). At OD_600nm_=0.4-0.5, cells were shifted from 37 °C to 16 °C and supplemented with 1 mM Bpa (dissolved in 1M NaOH; Bachem) and 100 µM ZnCl_2_. After 30 minutes, protein expression was induced by addition of 0.02% L-arabinose and 0.2 mM IPTG for ∼16 hours at 16 °C. Proteins were purified as described for the BRCA1^RING^/BARD1^216^ construct. Truncations that did not incorporate Bpa were separated via the SP column.

*BRCA1/BARD1 from insect cells:* For insect cell expression, truncation and deletion constructs were derived from full-length BRCA1 (pFastBac-FLAG-BRCA1^1-1863^; gift from W. Zhao, UT Health Sciences Center at San Antonio) and insect cell codon-optimized full-length BARD1 (pFastBac-Twin-StrepTagII-BARD1^1-777^; gift from A. Deans, Addgene plasmid #137166)(Tan et al., 2020) using Gibson assembly cloning (NEB). Baculovirus was generated using the Bac-to-Bac system in suspension culture *Sf9* cells according to the manufacturers protocols (Invitrogen). For general *Sf9* growth and virus amplification, SF900-II media (ThermoFisher) was supplemented with 5% FBS (HyClone) and 1x antibiotic-antimycotic (Gibco). For protein expression, 15 mL of BRCA1- and 15 mL of BARD1-containing P3 baculoviruses were added to 650 mL of *Sf9* cells at ∼1.5x10^6^ cells/mL in SF900-II media supplemented with 1% FBS and 1x antibiotic-antimycotic. Protein expression was carried out for 48-72 hours at 27 °C, shaking at 110 RPM. Following expression, cells were spun down at 150 x g for 10 minutes. The pellet was resuspended in 30 mL of Strep-start buffer (50 mM Tris-HCl pH 8, 10% glycerol, 150 mM NaCl, 1 mM DTT) supplemented with EDTA-free protease inhibitors and 1 mM PMSF and flash-frozen. Following freeze-thawing in a 37 °C water bath, additional EDTA-free protease inhibitors and PMSF were added in addition to 2 µL of benzonase (Sigma Aldrich). All subsequent steps were performed on ice or in a cold room over a short timeframe to prevent proteolysis (<3 hours total). Lysis was performed by sonication using a Branson Sonifier 250 (VWR Scientific) by applying 2 rounds of 15 pulses at 70% duty on power level 8, ensuring that the cell mixture stayed cold. PMSF was added throughout the lysis protocol. The lysed cells were centrifuged at 41,000 x g for 15 minutes, filtered through a 0.45 mM low protein-binding PVDF filter (Millipore) and applied to a 1 mL StrepTrap HP column pre-equilibrated in Strep-start buffer (Cytiva). The column was washed with 8 mL Strep-start buffer supplemented with protease inhibitors, 8 mL wash buffer (25 mM Tris-HCl pH 7.5, 300 mM NaCl, 0.01% Igepal-CA630, 1 mM DTT, 5 mM MgCl2, 2 mM ATP, 10% glycerol, EDTA-free protease inhibitors), and an additional 5 mL of Strep-start buffer. The column was eluted using 6 mL of Strep-start buffer supplemented with 3 mM d-Desthiobiotin (Sigma). The elution was diluted with 20 mL of ion-exchange buffer (25 mM HEPES-NaOH pH 7.5, 10% glycerol, 1 mM DTT) and applied to a 1 mL SP HP column pre-equilibrated in ion-exchange buffer containing 75 mM NaCl. The SP column was eluted over 12- CV (75-500 mM NaCl). Peak fractions were combined and concentrated using a 30K MWCO concentrator (Amicon), and flash-frozen in small aliquots. Concentrations were estimated by measuring absorbance at 280 nM using molar extinction coefficients obtained from ProtParam (Expasy). Concentration estimates were verified by SDS-PAGE and Coomassie staining, and slight adjustments in concentration were applied to best equalize the amounts of E3 in activity and binding assays.

*BARD1 intrinsically disordered region:* BARD1 124-270, 141-216, 269-424, and 124-424 were sub-cloned from pFastBac-6xHis-BARD1^1-777^ (gift from W. Zhao, UTHCSCA) into a modified pET28 vector containing an N-terminal 6xHis-SUMO tag using Gibson assembly cloning.

Plasmids were transformed into BL21 (DE3) *E. coli*, grown at 37 °C in M9 media supplemented with appropriate combination of isotopes (^15^N-NH_4_Cl, ^13^C-glucose; Cambridge Isotopes) to an OD_600nm_ of 0.6-0.8, and induced with 0.2 mM IPTG for ∼16 hours at 16 °C. HisTrap Ni^2+^ purification was performed as described above for BRCA1/BARD1 constructs expressed in *E. coli*. Elutions were dialyzed into ion exchange buffer (25 mM Tris pH 8.0, 200 mM NaCl, 0.5 mM EDTA, 1 mM DTT) in the presence of GST-SENP1 (produced in-house) to cleave the SUMO tag, leaving a non-native serine residue preceding the N-terminus except in the case of 269-424. The cleaved product was applied at a HiTrap Q column preceding a HiTrap SP column in- tandem to capture the non-specifically bound DNA (Cytiva). After application, the Q column was removed, and the SP column was eluted using a 12-CV gradient (0.2-1M NaCl). The peak fractions were combined, concentrated, and further purified using a Superdex 75 column equilibrated in NMR buffer (25 mM MOPS-NaOH pH 7.0, 100 mM NaCl, 1 mM EDTA, 0.5 mM TCEP (tris(2• carboxyethyl)phosphine)).

*Histone purification*: Wild-type and mutant human histones were transformed in *E. coli* BL21 (DE3) *pLysS* cells, grown at 37 °C to an OD_600nm_ of 0.5-0.6, and induced with 0.5 mM IPTG for 2 hours (H4) or 3 hours (H2A-2A, H2B-1K, H3.2) at 37 °C. Histone octamers used for fluorescence labeling were purified using the one-pot refolding protocol as described previously using 6xHis-TEV-H2A(Lee et al., 2015; Witus, Burrell, et al., 2021). For nucleosome ubiquitylation assays, a glycine within the TEV cleavage scar was mutated to a cysteine for fluorophore conjugation (H2A G-1C) via site-directed mutagenesis. For fluorescence based binding assays, a cysteine was introduced at H2B D51 via site-directed mutagenesis.

For reconstituting chemically modified H2A-Ub nucleosomes, histones were purified individually essentially as described(Luger et al., 1999). Briefly, inclusion bodies from 6-8L of histone expression in LB media were lysed in buffer T (50 mM Tris-HCl pH 7.4, 1 mM EDTA, 100 mM NaCl), washed by sonication twice in buffer TW (50 mM Tris-HCl pH 7.4, 1 mM EDTA, 100 mM NaCl, 1% v/v Triton-X100), and then twice again in buffer T. Inclusion bodies were softened by stirring with a small volume of DMSO (2-3 mL), and extracted with 60 mL of extraction buffer (25 mM Tris-HCl pH 7.5, 7 M guanidinium-HCl, 0.5 mM EDTA, 10 mM DTT). After centrifugation, the supernatant was dialyzed overnight into ion-exchange buffer (25 mM Tris-HCl pH 7.5, 7 M deionized urea, 100 mM NaCl, 0.5 mM EDTA, 10 mM DTT), and applied to 2x5 mL HiTrap Q columns preceding 2x5 mL HiTrap SP columns in-tandem to capture the non-specifically bound DNA (Cytiva). After sample application, the Q columns were removed, and the SP columns were eluted using a 12-CV salt gradient (0.1-1M NaCl for H2A and H2B, 0.2-1M NaCl for H3 and H4). The peak fractions were assessed by Coomassie stained SDS-PAGE gel and Abs_260_/Abs_280_ value, and the purest fractions were extensively dialyzed into water, aliquoted, flash frozen, and lyophilized. Lyophilized histones were reconstituted into H2A/H2B dimers, H3/H4 tetramers, and octamers as previously described(Dyer et al., 2003), and purified via SEC using Superdex 75 (H2A/H2B) or Superdex 200 (H3/H4 and octamers) columns in SEC buffer (25 mM Tris-HCl pH 7.5, 2M NaCl, 1 mM DTT).

*Fluorophore conjugation to histones and ubiquitin*: For fluorophore conjugation, Alexa Fluor 680 C_2_ maleimide (ThermoFisher), IRDye 680LT maleimide (Li-Cor), and Oregon green 488 maleimide (AAT Bioquest) were reconstituted in DMSO to 10 mM. Fluorophore maleimides were mixed with octamers containing single cysteine mutants H2A G-1C for H2A-Ub activity assays (with Alexa Fluor 680 C_2_ maleimide or IRDye 680LT maleimide) or H2B D51C for binding assays (with Oregon green 488 maleimide). Conjugation reactions were performed using 20 μM octamer and 100 μM fluorophore overnight at 4 °C in conjugation buffer (25 mM Tris-HCl pH 7.5, 2M NaCl, 1 mM EDTA, 0.5 mM TCEP), and stopped by addition of 10 mM DTT and flash frozen. Excess fluorophore was removed after nucleosome reconstitution by SEC using a Superdex 200 increase 10/300 GL column or extensive buffer exchange in a 30K MWCO concentrator (Amicon). Typical labelling efficiency was ∼70%. For labelling of Ub Q2C, 100 μM Alexa Fluor 680 C_2_ maleimide was mixed with ∼1 mM of Ub Q2C in buffer (25 mM Tris-HCl pH 7.5, 1 mM EDTA, 0.5 mM TCEP) at 4 °C overnight. The reaction was quenched with 10 mM DTT, and buffer exchanged into reducing buffer (25 mM Tris-HCl pH 7.5, 1 mM EDTA, 10 mM DTT) in a concentrator to remove unconjugated dye, and stored at -80 °C. The fluorophore labelling efficiency of Ub was calculated to be ∼5%.

*Ubiquitin G76C and dichloroacetone crosslinking of H2A-Ub*: H2A K15C, K119C, and K127C were generated by site-directed mutagenesis and purified as described above for individual histones. His-TEV-ubiquitin G76C was a gift from C. Wolberger (Addgene plasmid #75299) and purified as previously described(Morgan et al., 2019; Morgan Michael T. et al., 2016).

Dichloroacetone crosslinking of Ub G76C to H2A single cysteine mutants was performed as previously described in detail (Morgan et al., 2019), with the exception that the final product containing Ub, Ub-Ub dimers, and H2A-Ub was not purified by RP-HPLC to remove unconjugated Ub. Instead, it was lyophilized, and mixed with a slight excess of H2B in unfolding buffer (25 mM Tris-HCl pH 7.5, 7 M guanidinium-HCl, 0.5 mM EDTA, 1 mM DTT), refolded into H2A/H2B dimers in refolding buffer (25 mM Tris-HCl pH 7.5, 2M NaCl, 1 mM DTT), and purified by SEC using a 24 mL Superdex 75 column (GE Healthcare) to separate the H2A-Ub from Ub species.

*‘601’ and MMTV DNA:* For large scale purification of 147-bp ‘601’ (gift from K. Luger, CU Boulder), 185-bp ‘601’ (gift from S. Tan, Penn State Univ.), MMTV (gift from G. Debelouchina, (UC San Diego), and NLE-trimer ‘601’ (for tri-NCP; gift from K. Luger) DNA, repeat plasmids from 8-12L of DH5a cells in LB media were subjected to alkaline lysis, phenol-chloroform extraction, PEG precipitation, EcoRV digestion (NEB), and ion exchange of the excised fragment by HiTrap DEAE (147-bp ‘601’, 185-bp ‘601’, and MMTV; Cytiva) or mono-Q (NLE- trimer ‘601’; GE Healthcare) according to established protocols(Dyer et al., 2003).

For UV-induced Bpa crosslinking and EMSA experiments, fluorophore labelled ‘601’ DNA was generated using large-scale PCR with Phusion polymerase (produced in-house) from a pGEM- 3z/601 plasmid containing one copy of ‘601’ DNA (gift from J. Widom, Addgene plasmid #26656)(Lowary & Widom, 1998) with 5’-IRdye700 labelled forward primers (IDT). Following PCR, the reactions were purified using a 1 mL mono-Q column equilibrated in ion-exchange buffer (25 mM Tris-HCl pH 8.0, 200 mM NaCl, 1 mM EDTA) over a 10-CV gradient (0.2-1M NaCl). Peak fractions containing purified 147- and 185-bp DNA fragments were dialyzed into storage buffer (10 mM Tris-HCl pH 8.0, 1 mM EDTA), concentrated, and stored at -20 °C. The primer sequences were as follows: 147-fwd: ctggagaatcccggtgccgagg; 147-rev: acaggatgtatatatctgacacg; 185-fwd: atccctatacgcggccgccctggagaatcccggtgccgagg; 185-rev: atcgctgttcaatacatgcacaggatgtatatatctgacacg.

For generation of DNA fragments with DraIII sticky ends for dinucleosome assembly, large scale PCR was performed as described for the fluorophore-labelled DNA fragments above. Primer sequences containing DraIII cut-sites were based off those used by Poepsel *et al*. to make dinucleosomes with 35-bp of linker DNA(Poepsel et al., 2018). Fragments used to reconstitute nucleosomes containing fluorophore-labelled H2A nucleosomes (H2A^observe^) were generated using the “147-fwd” primer (sequence above) and the di-NCP reverse primer sequence (taggtatcgtat<u>CACGGGGTG</u>agatcgctacaggatgtatatatctgacacg). The fragment for the H2A^silent^ nucleosomes were generated using the “147-rev” and the di-NCP-forward primer sequence (ctgacttattga<u>CACCCCGTG</u>atgctcgatactgtcatactggagaatcccggtgccgag). The DraIII sites in the primers are underlined and uppercase. Following PCR purification by 1 mL mono-Q, DNA fragments were dialyzed into TE/0.1 buffer (10 mM Tris-HCl, 0.1 mM EDTA) and digested with DraIII-HF in 1x CutSmart buffer for ∼20 hr at 37 °C (NEB). Fragments with DraIII sticky ends were re-purified by 1 mL mono-Q ion-exchange, dialyzed into storage buffer (10 mM Tris-HCl, 1 mM EDTA), concentrated, and stored at -20 °C.

*Nucleosome reconstitution:* mono-nucleosomes (NCPs) were reconstituted by the standard salt- dialysis method by mixing histone octamers (or H3/H4 tetramers and H2A/H2B dimers) with ‘601’ DNA in high salt buffer (25 mM Tris-HCl pH 7.5, 2M NaCl, 0.1 mM EDTA, 1 mM DTT) in a slide-a-lyzer mini dialysis unit (0.1 mL, ThermoFisher)(Dyer et al., 2003). Low-salt buffer (25 mM Tris-HCl pH 7.5, 0.1 mM EDTA, 1 mM DTT) was pumped in over ∼24 hours in a cold room with stirring. A final dialysis was performed into NCP storage buffer (25 mM HEPES-NaOH, 10 mM NaCl, 0.1 mM EDTA, 1 mM DTT). Fluorophore-labelled nucleosomes were further purified by SEC using a 24 mL Superdex 200 increase 10/300 column equilibrated in NCP storage buffer or by extensive buffer exchange in a concentrator to remove excess fluorophore. For tri-NCPs, salt dialysis was performed as described for mono-NCPs using ∼1.4x octamer to ‘601’ sites in the presence of 0.2 equivalents of MMTV competitor DNA. Following dialysis into TEK10 (10 mM Tris-HCl pH 7.5, 0.1 mM EDTA, 10 mM KCl), trimers were precipitated by adding 4 mM MgCl_2_, incubating on ice for 10 minutes and pelleted by centrifugation at 17,000 x g at 4 °C. The supernatant was discarded, and the pellet was gently resuspended in TEK10, and dialyzed overnight into NCP storage buffer. Tri-NCP assembly was verified by EcoRI digestion of sites located between nucleosome units.

For di-NCP assembly, mono-NCPs with DraIII sticky ends were assembled as described above. The strategy to make asymmetric di-NCPs was based on previously described methods(Dao et al., 2020; Poepsel et al., 2018). 100 nM of each NCP was mixed in T4 ligase buffer (50 mM Tris-HCl pH 7.5, 10 mM MgCl_2_, 1 mM ATP, 10 mM DTT) on ice. T4 ligase (NEB) was added to a final concentration of 20 U/µL, the reaction was incubated for 30 minutes at 16 °C, and dialyzed into TEK10 for 4 hours at 4 °C. Following dialysis, MgCl_2_ was added to a final concentration of 18 mM, incubated at room temperature for 15 minutes, and pelleted at 17,000 x g at 4 °C. The pellet containing di-NCPs was resuspended in 50 µL of TEK10 and dialyzed overnight into NCP storage buffer. All chromatin substrates were analyzed for quality on 5% polyacrylamide 0.5x TBE gels monitoring fluorescence and/or DNA-staining, ensuring minimal free DNA or contaminating species. Chromatin substrates were stored on ice for no longer than one month.

*Design and assembly of DNA competitor fragments:* single-stranded, double-stranded, bubble, frayed, and single-stranded overhang DNA fragments were assembled by annealing ssDNA oligos (IDT) in IDT duplex buffer (30 mM HEPES-NaOH pH 7.5, 100 mM potassium acetate). Oligo mixtures were heated to 95 °C and slowly cooled to 25 °C stepwise over 1 hour in a thermocycler. For NMR experiments, dsDNA and bubble-DNA were buffer exchanged into NMR buffer in a spin concentrator (Amicon). The sequences of oligos used are as follows. For ssDNA, dsDNA, bubbleDNA, and ssOverhang a common “bottom” oligo was used (ggtacacaattgcgctggtaccccaggcgtcgtagg). The oligos annealed to the “bottom” oligo were as follows: (dsDNA-top: cctacgacgcctggggtaccagcgcaattgtgtacc; bubbleDNA-top: cctacgacgcctggctctttcccgcaattgtgtacc; ssOverhang-top: ccgcaattgtgtac). For the frayed DNA, the annealed oligos were as follows: (frayedDNA-top: ggtacacaattgcgccaggcgtcgtaggctggtacc; frayedDNA-bottom: ctctttcccctacgacgcctggcgcaattgtgtacc).

### Nucleosome ubiquitylation assays

*Time-course H2A-Ub assays:* The general setup for time-course H2A-Ub assays was a reaction mixture containing 0.2 µM E1 (UBA1), 1 µM E2 (UBE2D3), 30 µM ubiquitin, 0.5 µM NCP substrate, and 12.5-100 nM E3 (depending on the assay), assembled in 30 µL of reaction buffer (25 mM HEPES-NaOH pH 7.5, 150 mM NaCl). For assays containing di-NCPs, 10 nM of fluorophore-labelled di-NCP substrate was mixed with 0.5 µM unlabeled NCP^147^ substrate in the same reaction tube to facilitate quantifiable reaction kinetics. Critically, the salt concentration of each reaction within an assay was rigorously controlled, as H2A-Ub kinetics are extremely sensitive to differences in buffer ionic strength. 5 µL of each reaction was mixed 1:1 with 2x SDS-PAGE load dye as a zero time-point. The mixtures were brought to 37 °C in a heat block, and 1 µL of 100 mM MgCl_2_/ATP was added (4 mM final concentration) to initiate the reaction, and the indicated time-points were obtained by diluting 5 µL the reaction 1:1 with 2x SDS-PAGE load dye. For experiments with tri-NCPs, the concentration of nucleosome units in the reaction was normalized to NCP^147^ and NCP^185^ (0.5 µM NCP unit or 0.17 µM tri-NCP), and reactions were initiated with 2 mM ATP and 1 mM MgCl_2_.

Gel samples were run on 15% SDS-PAGE gels and visualized by in-gel fluorescence monitoring a fluorophore conjugated to the N-terminus of H2A. Alternatively, some assays were visualized by Western blot for H2A (EMD Millipore, 07-146) or a VSVG epitope tag on the N-terminal tag of H2A (Sigma, V4888) and a fluorescently labelled secondary antibody (Cell Signaling, 5151S).

Gels and blots were imaged on a Li-Cor Odyssey imager (Li-Cor). Band quantification was performed using Image Studio software (Li-Cor) by drawing a box encompassing the unmodified H2A band to measure its intensity, performing background subtraction, and normalizing to the zero time-point of a given reaction.

*Inhibition of H2A-Ub by competitor DNA:* A reaction mixture containing 0.2 µM E1 (UBA1), 3 µM E2 (UBE2D3), 30 µM ubiquitin, 0.5 µM NCP substrate, and 50 nM E3 (or 750 nM for BRCA1^RING^/BARD1^RING^) was assembled in 9 µL of reaction buffer (25 mM HEPES-NaOH pH 7.5, 100 mM NaCl) in PCR strips. To that, 1 µL of DNA competitor at 10x concentration was added. Reactions were brought to 37 °C in a thermocycler, started by addition of 5 µL of 10 mM ATP/MgCl_2_ in reaction buffer, and allowed to proceed for 12-20 minutes depending on the E3.

Reactions were stopped by addition of 10 µL of 4x fluorescence sample load dye (Li-Cor), resolved on a 15% SDS-PAGE gel, and visualized using a Li-Cor Odyssey imager.

Quantification was performed as described above, monitoring the intensity of unmodified H2A, and normalizing each reaction to a -ATP reaction.

*Auto-ubiquitylation assay:* Reactions were assembled with 0.2 µM E1 (UBA1), 1 µM E2 (UBE2D3), 10 µM ubiquitin (Q2C-Alexa Fluor 680), and 0.2 µM of the indicated E3 in buffer (25 mM HEPES-NaOH pH 7.5, 50 mM NaCl), and were initiated by addition of 5 mM ATP/MgCl_2_.

After 60 minutes at 37 °C, reactions were quenched by mixing 1:1 with 4x fluorescence sample load dye (Li-Cor), resolved on a 4-20% SDS-PAGE gel (BioRad), and visualized using a Li-Cor Odyssey imager.

### Nucleosome binding assays

*Electrophoretic mobility shift assay:* For EMSA experiments, 5 µL of 10 nM of substrate (NCPs with either H2A G-1C labelled with Alexa Fluor 680 or DNA/NCPs labelled with IRDye700) were mixed with 5 µL of indicated RING/RING heterodimer or BARD1 IDR constructs (2x concentrated) in EMSA buffer (15 mM Tris-HCl pH 7.5, 90 mM NaCl, 1 mM DTT, 0.1 mg/mL BSA, 0.05% triton X-100) in PCR strips. Binding was allowed to proceed for 10 minutes at room temperature, followed by 5 minutes on ice. 5 µL of 25% sucrose was added to each tube, and 15 µL of sample was loaded onto 5% polyacrylamide native gels (0.5x TBE) in pre-chilled 0.5x TBE buffer. Gels were run at 4 °C for 90-120 minutes at 110V and visualized using a Li-Cor Odessey scanner (Li-Cor).

*Fluorescence-quenching binding experiments:* Assay set up was performed essentially as previously described (Hu et al., 2021). 10 μL of 10 nM NCPs with Oregon green 488 conjugated to H2B D51C in low-salt fluorescence buffer (25 mM Tris-HCl pH 7.5, 10 mM NaCl, 0.01% CHAPS, 0.01% NP40, 0.1 mg/ml BSA, 1 mM DTT) were mixed with the 10 μL of the indicated BRCA1/BARD1 construct at 2x concentration in high-salt fluorescence buffer containing either 90 mM, 190 mM, or 290 mM NaCl, depending on the assay, in a 384-well low volume black round bottom polystyrene non-binding surface microplate (Corning). Plates were briefly mixed, incubated at 22 °C for 10-15 minutes, and scanned using a BioTek Synergy Neo2 plate reader equipped with a fluorescence filter (excitation: 485/20 nm, emission: 528/20 nm). The gain was set to 100 for all experiments. Two to three technical replicates for each independent replicate experiment (fresh NCP and BRCA1/BARD1 dilutions) were performed, and the readings from two scans were averaged. BRCA1/BARD1/nucleosome binding interactions were quantitatively analyzed by fluorescence quenching assuming the following systems of equations:

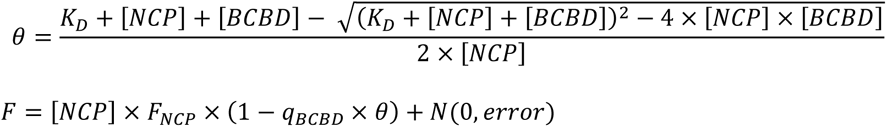

where θ is the fraction of nucleosome bound by BRCA1/BARD1 (BCBD), F_NCP_ is the fluorescence per nucleosome, q_BCBD_ is the fraction of fluorescence quenching per BRCA1/BARD1 binding, and fluorescence measurements are gaussian distributed with mean 0. Maximum likelihood parameter estimates were determined using custom Python scripts with optimization by a differential evolution algorithm implemented in Scipy (Virtanen et al., 2020). K_d_ and q_BCBD_ were separately estimated for each BRCA1/BARD1 construct, and NCP concentration was treated as a random effect per individual dilutions from higher concentration stock solutions, with the total number of model parameters being equal to number of NCP dilutions + (2 x number of BCBD constructs) + 2. The 95% confidence limits were estimated by computing the profile likelihood with an applied Bonferroni correction.

### UV-induced Bpa crosslinking

NCPs and DNA for UV-induced crosslinking assays were reconstituted using a 185-bp ‘601’ sequence labeled with an IRdye700 fluorophore (described above). 50 nM NCPs or DNA were incubated with 500 nM of the BRCA1^1-104^-(GS)_6_-BARD1^26-221^ fusion (wild-type or with different Bpa-incorporated sites L120Bpa, W146Bpa, Y180Bpa, A195Bpa, W218Bpa) for 15 minutes at 4°C in 30 uL UV Crosslinking buffer (20 mM HEPES-NaOH pH 7.5, 100 mM NaCl) in a 96-well clear round bottom polystyrene non-treated microplate (Corning). Following incubation, 5 µL was added to 5 µL 2X SDS-PAGE gel-loading dye for the -UV sample. Samples were then subjected to UV irradiation (365 nm) using a Blak-Ray B-100AP/R High Intensity UV Lamp (UVP) from 20 cm for 30 minutes at 4°C. Following irradiation, 5 µL was added to 5 µL of 2x SDS-PAGE gel-loading dye. Samples were heated at 60°C for 5 minutes and separated on 10% SDS-PAGE gels. Gels were visualized using a Li-Cor Odyssey scanner (Li-Cor). Band quantification was performed using Image Studio software (Li-Cor) by drawing a box measuring the total band intensity for each crosslinked product, subtracting the background signal, and normalizing to the crosslinked band of the L120Bpa fusion mutant. For the Bpa UV crosslinking DNA competition assays, 50 nM NCPs were incubated with 500 nM BRCA1-f-BARD1^221^ (W146Bpa or A195Bpa) with varying amounts of competitor 36-mer dsDNA or bubble-DNA (0, 0.3125, 0.625, 1.25, 2.5, 5, or 10 μM) for 15 minutes at 4°C, and assays were performed as described above.

### Cellular Studies

*Mammalian cell culture and transfection:* HeLa cells were grown in Dulbecco’s modified Eagles medium (DMEM) supplemented with 10% fetal bovine serum (Sigma), 100 μg/ml streptomycin, and 100 U/ml penicillin (Sigma). The cells were tested for mycoplasma contamination by Bionique testing labs (http://www.bionique.com/). To create a FLP-in version of HeLa, we stably integrated a flippase recognition target (FRT) sequence into the cells by using the pFRT/lacZeo plasmid (Thermo Fisher Scientific). We tested Zeocin-resistant clones that had a single integration site detected by Southern blot for high-activity integration sites by using the mammalian b-galactosidase activity assay (Gal-Screen, Thermo Fisher Scientific). Clonal expansion of the selected colony established the HeLa-FRT cell line. To generate stable HeLa-FRT shBARD1 cells, Dharmacon™ TRIPZ™ lentiviral shRNAs against BARD1 (RHS4696) were transfected with their respective plasmids and individual clones were selected with 2 μg/ml puromycin. To generate HeLa-FRT shBARD1 cell lines expressing HA-BARD1 WT or mutant (Δ194-216), cells were transfected with their respective plasmids and individual clones were selected with 200 μg/ml hygromycin. BARD1 was cloned into pcDNA5/FRT-HA vector.

QuikChange site-directed mutagenesis was used to construct the mutant variant (Δ194-216) of BARD1.

*Immunoblot analysis*: HeLa-shBARD1 cells stably expressing wild type or Δ194-216 mutant of HA-BARD1 were pretreated with doxycycline(1μg/ml) for three days to knockdown the endogenous BARD1, and then incubated with or without Olaparib (10 μM) for another 24 h.

Cells were trypsinized and pellets were washed by PBS. Protein was extracted using NETN buffer (20 mM Tris-HCl pH 8, 420 mM NaCl, 1 mM EDTA, 0.5% IGEPAL, 1 mM DTT, and Roche Protease Inhibitor Cocktail) and were resolved in 8% Bis-Tris SDS-PAGE gels, and blots (20-50 µg of total protein) were probed with the following antibodies: HA (3724S, Cell Signaling; 1:1500), BRCA1 (SC6954, Santa Cruz; 1:500), BARD1 (ab50984, Abcam; 1:1000), Tubulin (2128S, Cell Signaling; 1:2000), Lamin B1 (SC374015, Santa Cruz, 1:500), according to the instructions provided by the manufacturers. The blots were incubated with HRP-conjugated secondary antibodies (Pierce 31450 for rabbit anti-mouse IgG-HRP; Sigma A6154 for goat anti-rabbit IgG-HRP) before being visualization of protein signals using the ECL max kit (Bio-rad).

*Preparation of cytoplasmic and nuclear extracts:* The REAP method for the preparation of cytoplasmic and nuclear extracts was followed. Briefly, HeLa cells from 10-cm dishes were washed with ice-cold phosphate buffer saline (PBS) pH 7.4, collected by centrifugation, resuspended in 900 μL of ice-cold PBS with 0.1% NP40 and protease inhibitors, and triturated 5 times using a p1000 micropipette. The lysed cell suspension was centrifuged for supernatant (this is the cytoplasmic fraction) and the pelleted nuclei was washed once with PBS with 0.1% NP40 and lysed by NETN buffer with protease inhibitors and different NaCl concentrations (100, 300 and 420 mM) to yield the nuclear extract fractions. Each soluble fraction was labeled as NS100, NS300 and NS420, while the final pellets were resuspended with NETN420 buffer and sonicated for 10 seconds for NP420 fraction. The cytoplasmic and nuclear fractions, 20 µg each, were analyzed by immunoblotting for their content of BRCA1, BARD1, Tubulin and Lamin B1.

*Immunofluorescence microscopy and image analysis:* Cells were pretreated with doxycycline (1μg/ml) for three days to knockdown the endogenous BARD1. Cells were trypsinized and 4x10**^4^** cells seeded in the 4-chambers slide with or without Olaparib (10 μM) for another 24 h. Cells were pre-extracted on ice for 10 min with cold cytoskeleton buffer (10 mM PIPES pH 6.8, 100 mM NaCl, 300 mM sucrose, 3 mM MgCl2, 1 mM EGTA, 0.5% Triton X-100) followed by 10 min with cytoskeleton stripping buffer (10 mM Tris-HCl pH 7.4, 10 mM NaCl, 3 mM MgCl2, 1% Tween 20 (v/v), 0.5% sodium deoxycholate). Pre-extracted cells were then fixed at room temperature for 10 min with 4% paraformaldehyde, washed with PBS, and treated for 10 min with 1% Triton X-100 in PBS. Primary anti-HA antibody (3724S, Cell Signaling; 1:500) was incubated overnight at 4°C in 5% goat serum in PBS. Alexa Fluor-568 conjugated secondary antibody (Jackson Immunno Research Labs) was incubated for 1 h at room temperature, and DAPI were incubated for another 15 mins in room temperature. Then slides mounted using antifade mounting media (9071, Cell signaling). A random selection of 10-11 areas were imaged using an Evos M5000 Microscope. HA-BARD1 foci>5 and HA-BARD1 foci per nuclear were quantified by the Celleste software and Prism GraphPad software was used for the final figure and significance analysis.

*Clonogenic survival assay:* HeLa cell lines stably expressing HA-BARD1 wild type or mutant were pretreated with doxycycline for 3 days. Then, 400 cells/well were seeded into 12-well plates, treated with indicated amount of Olaparib (Selleckchem) or Cisplatin (Selleckchem) in regular growth medium for 11-12 days. Cells were fixed with methanol and stained with 0.5% crystal violet in methanol before colonies were counted. Clonogenic survival was determined for a given concentration of cells that were plated by dividing the number of colonies on each treated plate by the number of colonies on the untreated plate, taking the plating efficiency of untreated cells into account.

### Chemical Crosslinking and Mass Spectrometry Analysis

*Sample preparation:* Reactions were 90 µL of protein/nucleosome mix containing 2 mM BRCA1^RING^/BARD1^FL^ and 2 mM of unmodified H2A or H2A K15-Ub nucleosomes wrapped with 147-bp ‘601’ DNA in crosslinking bufffer (25 mM HEPES-NaOH pH 7.5, 150 mM NaCl, 1% glycerol, 1 mM DTT) plus 3.06 µL of 14.5 mM DSS or BS3 (final concentration 0.5 mM) (DSS: disuccinimidyl suberate, BS3: bis(sulfosuccinimidyl)suberate, ThermoFisher). Crosslinking was carried out for 10, 30 or 45 mins at room temperature before removing 30 µL and quenching by addition of 3 µL of 1 M ammonium bicarbonate at room temperature for 30 minutes. Four separate reaction conditions were performed: (1) WT-NCP DSS; (2) WT-NCP BS3; (3) K15Ub- NCP DSS; (4) K15Ub-NCP BS3. After quenching, 9 µL of each sample at each timepoint was analyzed by SDS-PAGE **(see Extended Data Fig 10a)** and the remainder was stored at -80°C until processing for mass spectrometry analysis.

Reactions were prepared for MS analysis by bringing them up to 0.1% PPS silent surfactant (Expedion Inc.), 5 mM TCEP. Samples were reduced for 60 mins at 60°C in an Eppendorf Thermomixer with shaking (1200 rpm). Alkylation was performed at room temperature in the dark for 30 mins with 6 mM iodoacetamide. Excess iodoacetamide was quenched by addition of 5 mM DTT and samples were then digested by trypsin digestion (Sequencing Grade Modified Trypsin, Promega Corp) at 37°C for 6 hours in a Thermomixer with shaking (1000 rpm) at a substrate to enzyme ratio of 15:1 prior to acidification with 250 mM HCl (final concentration).

Acidified samples were incubated for 1 hour at room temperature, spun at max speed in a microfuge and supernatants containing peptides were transferred to autosampler vials and stored at •80°C until analysis.

*Chromatography:* Mass spectrometry and data analysis was based on previously described methods(Zelter et al., 2015). For each injection, 3 µl of protein digests were loaded by autosampler onto a 150­μm Kasil fritted trap packed with 2 cm of Reprosil­Pur C18­AQ (3­μm bead diameter, Dr. Maisch) at a flow rate of 2.5 µl/min. After desalting with 8 µl of 0.1% formic acid plus 2% acetonitrile, the trap was brought online with a Self-Packed PicoFrit Column (New Objective part number PF360-75-10-N-5, 75 µm i.d.) packed with 30 cm of Reprosil•Pur C18­AQ (3­μm bead diameter, Dr. Maisch) mounted to a heated nanospray ionization source (CorSolutions LLC) set at 50°C and placed in line with a Thermo Scientific EASY-nLC 1200 UPLC pump plus autosampler.

Peptides were eluted from the column at 0.25 µL/min using an acetonitrile gradient consisting of the following steps: (1) 0-10 mins; 6-10% B; (2) 10-90 mins; 10-32% B; (3) 100-130 mins; 32-75% B; (4) 130-135 mins; 75% B; (5) 135-136 mins; 75-100% B; (6) 136-151 mins; 100% B, followed by re-equilibration to 0% buffer B prior to the subsequent injection. Buffer A was: 0.1% formic acid in water and buffer B was 0.1% formic acid 80% acetonitrile.

*Data acquisition:* A Thermo Fisher Scientific Exploris 480 was used to perform mass spectrometry in positive ion mode with the following settings. Data dependent acquisition (DDA) mode was used with a maximum of 20 tandem MS (MS/MS) spectra acquired per MS spectrum (scan range of *m/z* 400–1,600). The resolution for MS and MS/MS was 60,000 at *m/z* 200. The normalized automatic gain control targets for both MS and MS/MS were set to 100%, and the maximum injection times were 50 and 100 ms for MS and MS/MS scans, respectively. MS/MS spectra were acquired using an isolation width of 2 *m/z* and a normalized collision energy of 27. MS/MS acquisitions were prevented for +1, +2, ≥+6 or undefined precursor charge states.

Dynamic exclusion was set for 10 seconds. MS and MS/MS spectra were collected in centroid mode.

*MS data processing – identification of crosslinked peptides:* Raw mass spectra were converted into mzML using ProteoWizard’s msconvert(Chambers et al., 2012). All proteins in the sample were identified using Comet(Eng et al., 2013) searching against a database including the entire *S. frugiperda*, *E. coli* and *H. sapiens* proteomes (Uniprot 7108, 83333 and 9606, respectively) plus common contaminants (https://www.thegpm.org/crap/) along with the sequences of all heterologously expressed proteins (FLAG-BRCA1, P38398; TwinStrepII-BARD1, Q99728; H2A- 2A, Q99728; H2B-1K, O60814 ; H3.2 (C110A), Q71DI3; H4, P62805; Ub G76C, sequence derived from P0CG48). Smaller search databases were made for subsequent XL searching consisting only of proteins identified in initial comet searches by at least 3 peptides with a Percolator assigned q-value of ≤ 0.01. Decoy databases consisted of the corresponding set of reversed protein sequences. Cross•linked peptides were identified within these proteins by Kojak version 2.0.0-alpha8 available at (http://www.kojak-ms.org/)(Hoopmann et al., 2015). A statistically meaningful q•value was assigned to each peptide spectrum match (PSM) through analysis of the target and decoy PSM distributions using Percolator version 2.08(Käll et al., 2007). All data reported in this paper were filtered to show hits to the target proteins that had a Percolator assigned peptide level q value ≤ 0.05.

### NMR experiments and resonance assignments

All spectra were collected at 25° C in NMR buffer (25 mM MOPS-NaOH, 100 mM NaCl, 1 mM EDTA, and 0.5 mM TCEP, pH 7.0) with 7% D_2_O. NMR HSQC and titration data were completed on a Bruker 800 MHz Avance III spectrometer with cyroprobe. (^1^H,^15^N)-HSQC spectra were collected on the following ^15^N-BARD1 IDR constructs: BARD1 Ser-124-424, Ser-124-270, 269- 424, and Ser-141-216.

(^1^H,^15^N)- and (^1^H,^13^C)-HSQC titration experiments were completed for BARD1 IDR + 36mer- dsDNA or 36mer-bubDNA by adding unlabeled DNA to 150 µM [^13^C,^15^N]-BARD1 124-270 or 150 μM [^13^C,^15^N]-BARD1 Ser-141-216, maintaining a constant concentration of the labeled species. Only apo and 1:1 spectra for the [^13^C,^15^N] BARD1 Ser-141-216 construct were obtained since intermediate titration points resulted in loss of sample. Additionally, titration spectra with nucleosome were collected for 150 μM [^13^C,^15^N]-BARD1 124-270 + 0.05x NCP^147^. A (^1^H,^15^N)-HSQC was collected on 150 μM ^15^N-BARD1 269-424 + 150 μM 28mer-dsDNA.

Dynamics experiments were collected on a Bruker 500 MHz Avance III spectrometer with a room temperature probe. T_1_ and T_2_ ^15^N-Trosy HSQC experiments for 150 μM [^13^C,^15^N]-BARD1 Ser-141-216, 150 μM [^13^C,^15^N]-BARD1 Ser-141-216 + 150 μM 36mer-dsDNA, and 150 μM [^13^C,^15^N]-BARD1 Ser-141-216 + 150 μM 36mer-bubble-DNA were collected in an interleaved manner with 8 points each(G. Zhu et al., 2000). T_1_ delays were 10, 40, 80, 120, 160, 320, 640, and 1000 ms; T_2_ cpmg loop delays were 8.48, 16.96, 25.44, 25.44, 42.40, 50.88, 67.84, and 84.80 ms. T_1_ and T_2_ for each residue were fitted to a single exponential with errors reflecting the quality in the fit. Residues corresponding to peaks with overlapping intensities were excluded from the analysis.

Backbone chemical shift assignments were determined from standard backbone triple-resonance experiments (HNCA, HNCOCA, HNCOCACB, HNCACB, HNCO, and HNCACO) and an HNHA experiment using a Bruker 800 MHz Avance III spectrometer with cryoprobe. The 3D data were collected for 500 μM [^13^C,^15^N]-BARD1 Ser-124-270 and 500 μM [^13^C,^15^N]-BARD1 Ser-141-216. Assignments from the shorter construct were transferred as appropriate to the longer BARD1 construct. HNCACB and HNCOCACB spectra were collected on the 1:1 samples of 150 μM [^13^C,^15^N]-BARD1 Ser-141-216 with either 36mer dsDNA or 36mer bubble-DNA.

The NH chemical shift perturbation was calculated according to Δδ_NH_ (ppm) = sqrt [Δδ_H_ + (Δδ_N_/5)^2^]. Component ^1^H and ^15^N chemical shift differences are (free – bound) in ppm. The CaCb CSP was calculated according to Δδ_CaCb_ (ppm) = 0.25*[(Cα-Cβ)_free_-(Cα-Cβ)_bound_].

Secondary structure propensity (SSP) was determined from Δδ(Cα-Cβ) = (Cα-Cβ)_measured_ - (Cα- Cβ)_random coil_, where the random coil shifts were generated from BARD1 IDR sequence using a webserver (https://spin.niddk.nih.gov/bax/nmrserver/Poulsen_rc_CS/)(Kjaergaard & Poulsen, 2011).

### Multiple Sequence Alignment

Mammalian BARD1 ortholog sequences were downloaded from Ensembl (v106), globally aligned using MAFFT(Katoh & Standley, 2013), and trimmed to BARD1 residues 124-270.

### Materials, Data, and Code availability Statement

Source data for H2A-Ub and binding assays as well as uncropped gels and blots and original microscopy images are included as supplemental files. For crosslinking mass-spectrometry analysis, the complete unfiltered list of all PSMs and their Percolator assigned q values are available on the ProXL web application(Riffle et al., 2016, 2019) https://proxl.yeastrc.org/proxl/p/bard1nucleosome along with the raw MS spectra and search parameters used. In addition, complete search algorithm configuration files, fasta search databases, raw search output and raw MS data files were deposited to the ProteomeXchange Consortium via the PRIDE(Vizcaíno et al., 2009) partner repository with the dataset identifier PXD035345. NMR chemical shift assignments for BARD1 Ser-141-216 and associated DNA- bound assignments have been deposited to the Biological Magnetic Resonance Bank with dataset identifiers 51523 (apo), 51524 (with dsDNA), 51525 (with bubble-DNA). Python scripts used to analyze and plot fluorescence-based binding data are included with this manuscript.

Plasmid reagents generated in this study can be obtained upon request from the corresponding author Dr. Rachel Klevit.

